# Corepressor NCoR1-mediated regulation of mucin dynamics governs gut inflammation

**DOI:** 10.64898/2026.05.02.722388

**Authors:** Yesheswini Rajendran, Bhagyashree Srivastava, Preksha Gaur, Rohan Babar, Neha Guliya, Aamir Suhail, Lalita Mehra, Muskaan Kalra, Mukesh Singh, Prasenjit Das, Vineet Ahuja, Chittur V. Srikanth

**Affiliations:** Laboratory of Gut Infection and Inflammation Biology, Regional Centre for Biotechnology, 3^rd^ milestone, Faridabad-Gurugram expressway, Faridabad. India 121001; Harvard Medical School, USA; All India Institute of Medical Sciences, New Delhi, India; Bennett University, India

**Keywords:** NCoR1, KLF16, Inflammatory Bowel Disease (IBD), mTOR, Goblet cells, MUC2

## Abstract

Inflammatory bowel disease (IBD), comprising Ulcerative colitis (UC) and Crohn’s Disease, is a chronic relapsing immune-mediated inflammatory disorder of the gut. The intestinal mucus layer is a protective barrier that safeguards direct exposure of epithelium to luminal microbes and antigens. A prolonged disruption of the mucus layer may contribute to the development of IBD. Loss of mucin-producing goblet cells is a hallmark of UC. The underlying molecular mechanism controlling goblet regulation remains poorly understood. In the current work, we show a key role for NCoR1 (Nuclear corepressor 1) in goblet cell regulation. A specific downregulation of NCoR1 in intestinal crypts and goblet cells was observed in human UC and mice models. While NCoR1 was upregulated during goblet cell differentiation, inflammatory cues downregulated its expression. Experimental loss of NCoR1 resulted in exacerbated disease in a murine model of colitis, whereas its upregulation via Vitamin D led to a rescue. ChIP-seq led to the identification of KLF-16, a transcription factor, as a target of NCoR1. NCoR1 -KLF16 regulatory axis regulated key goblet cell proteins, including MUC2. Mechanistically, the regulation of MUC2 is modulated by the NCoR1-KLF16 axis, via mTOR signalling. In conclusion, this work shows a critical involvement of NCoR1-KLF16 in governing goblet cell function and intestinal homeostasis.

## Introduction

Inflammatory Bowel Disease (IBD) is a chronic, immune-hyperactive disease broadly classified into Ulcerative colitis (UC) and Crohn’s Disease (CD). The symptoms, diarrhoea, abdominal cramps, rectal bleeding, and weight loss manifest in cycles of remission and relapse, progressively leading to tissue damage and deleterious outcomes^123^. In case of UC, disease pathophysiology is limited to the mucosal layer and is an idiopathic inflammatory condition primarily restricted to the colon, causing superficial damage to the bowel wall^3^. Owing to a lack of cure, IBD patients live a very compromised lifestyle, and the disease poses a major health challenge^45^.

At the cellular level, IBD is due to failed relationship between gut epithelium and luminal microbes, leading to a heightened immune response. The exaggeration of immune response is triggered by dysregulated epithelial signalling, often arising due to altered microbial composition, defective antigen sensing, or mucus layer abnormality. In healthy tissue, luminal microbes are prevented from directly contacting host epithelium by a protective mucus layer. Loss of goblet cells (GCs), the cells responsible for the production of mucin, is a key hallmark of UC pathogenesis^6,7^.

GCs are specialised epithelial cells, originate from crypt resident stem cells, to secrete mucin (predominantly MUC2 in the colon) and maintain the mucus layer^8^. A healthy colonic epithelium is composed of 16% of GCs, which is significantly reduced in UC^9,10^. Also, GCs are involved in cytokine secretion and antigen presentation, thereby contributing to GCs - immunocyte cross-talk^11^. While, several transcription factors are involved in cell fate determination in the gut, the detailed mechanism of GCs differentiation remains poorly understood. Transcription factors, KLF4 and SPDEF are involved in goblet cell differentiation, mediating mucin MUC2 regulation^12^.

In a recent work, we showed the involvement of SENP7, a deSUMOylase enzyme, as a mediator of epithelial signalling in IBD. Enhanced SENP7 expression led to signalling events culminating in pathogenic gamma-delta γδT-cell infiltration and tissue inflammation. Interestingly, NCoR1, Nuclear Receptor Corepressor 1 protein, was among the several other interactors, identified to physically interact with SENP7^1^.

NCoR1 is a scaffolding protein involved in modulating gene repression via forming large protein complexes and recruitment of HDACs^13,14^. NCoR1 involvement in the maintenance of gut homeostasis has been cited for its importance in protection during DSS-mediated inflammation^15^; however, the mechanistic details remain unexplored. A balanced expression of NCoR1 in immune cells is essential for maintaining homeostasis, but NCoR1-mediated regulation in epithelial cells, in particular, GCs, during ulcerative colitis pathogenesis remains unexplored. Our data demonstrates that dysregulated NCoR1 in the colonic crypts harbouring goblet cells, dysregulates mucin release and affects goblet cell function, leading to exacerbated colitis.

## RESULTS

### Downregulated expression of NCoR1 in colon crypts in UC

To investigate the possible role of NCoR1 in UC pathogenesis, human UC endoscopic samples was analysed. To ensure statistical power >80%, a total of 79 human samples (33 non-IBD control subjects) and (46 UC), was included for different experiments described below. Compared to control, inflammation in the UC was evident by morphological and histopathological markers including immune infiltration, crypt abscesses (Fig.1a) and reduced number of GCs per crypt (Fig.S1a). Colonic biopsies of control and UC were sectioned and subjected to Immunohistochemistry (IHC) of NCoR1, followed by Alcian Blue-Nuclear fast red staining. Here, Alcian blue staining was done to locate GCs. NCoR1 expression was seen in the crypts of control, but drastically reduced in those areas of UC. In these UC sections, however, an increased expression of NCoR1 was seen in the infiltrated immune cells (Fig.1a). In UC, NCoR1expression was downregulated (∼-10-fold) seen by RT-PCR (Fig.1b) and similarly by immunoblot (Fig.1c). Taken together, these data reveal downregulation of NCoR1 in UC. Hence, understanding NCoR1 during colitis progression becomes important.

**Fig. 1:**
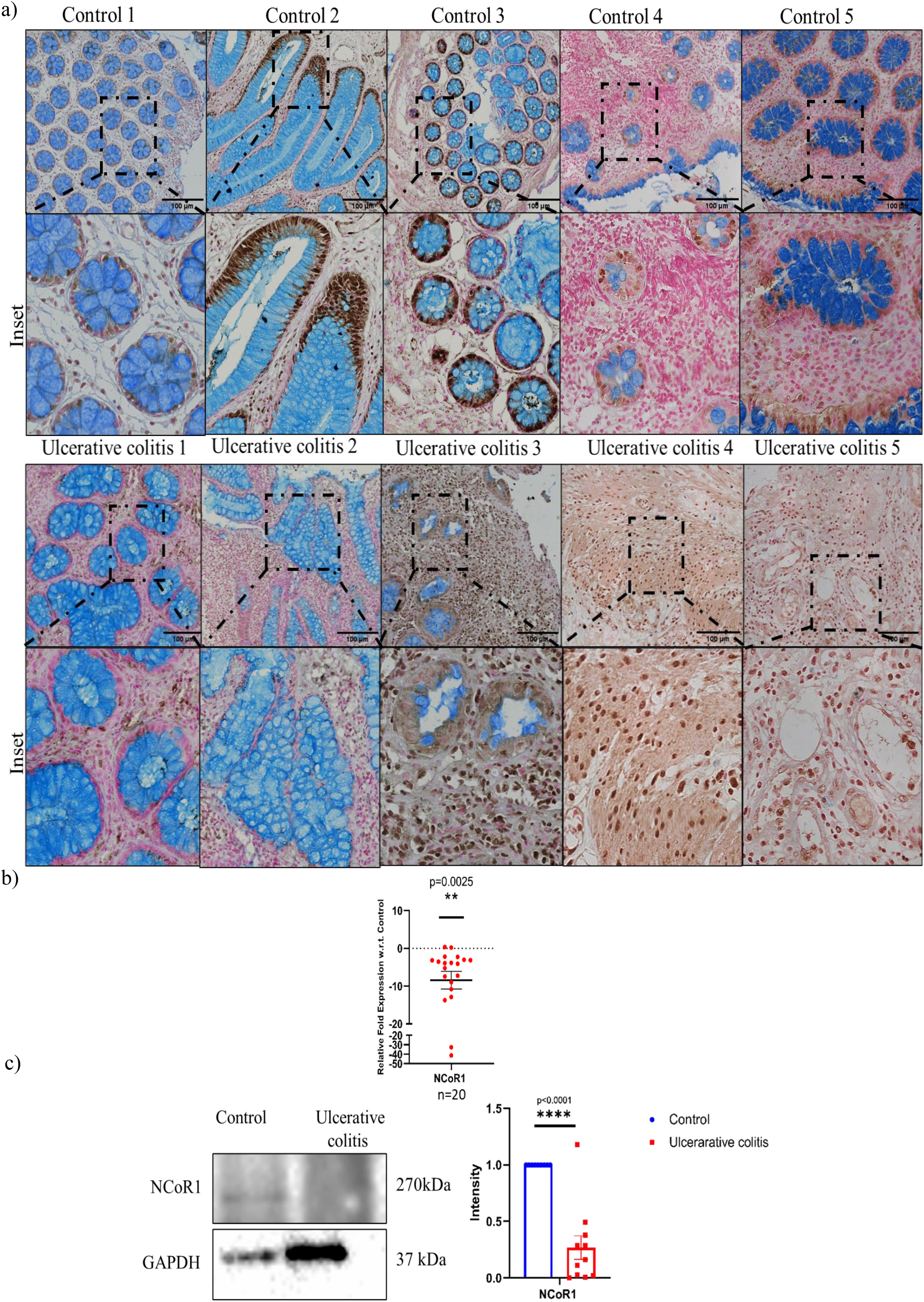
Expression of Corepressor NCoR1 in human ulcerative colitis. a) IHC-Alcian blue images showing NCoR1staining (brown colour) analysed in colon tissue biopsies sections of healthy individuals (n=5) and UC patients(n=5) (Scale bar= 100 µm). The inset shows zoomed-in areas of the image. Alcian blue (blue colour) represents colonic goblet cells and Nuclear fast red (red colour) represents nuclei. b) qRT-PCR analysis of relative fold expression of NCoR1 gene in human UC patient colonic biopsies (n=20) relative to average control value (n=24). 18s was used for normalisation. c) Representative immunoblot of NCoR1 protein in human ulcerative colitis (UC) (n=11) and control (n=9) biopsy samples. GAPDH was used as a loading control. Graph on the right showing densitometric analysis of NCoR1 expression normalised to control GAPDH. Each dot represents (B, C) one human. All data were expressed as means ± SEM, unpaired Student t-test was used to calculate statistical significance \**P* < 0.05; \*\**P* < 0.01; \*\*\**P* < 0.001; \*\*\*\**P* < 0.0001; ns, not significant. See also Fig S1 and Table S1.

### Altered expression of Corepressor NCoR1 in the colon in murine colitis

For detailed investigation of significance of NCoR1 in UC, a Dextran Sodium Sulphate (DSS) murine model of colitis was utilised^161^. The mice cohort were given 2% DSS in drinking water for different durations, i.e., 3 days (DSS-3), 5 days (DSS-5) and 7 days (DSS-7). Separately, one cohort of mice fed with only drinking water, was included as a control group. Body weight change was monitored daily (Fig 2b). Post euthanasia of the animals at the stipulated time, a significant alteration of colon morphology and reduction in length, was observed in DSS-5 and DSS-7 mice (Fig 2c) (Fig S2 a, b). Based on the various phenotype, DSS-1 and DSS-3 depict mild colitis, DSS-5 moderate colitis, and DSS-7 represents the severe colitis. Immunoblotting of colon lysates revealed a dynamic expression of NCoR1. In mild inflammation (DSS-3) NCoR1 was downregulated, but this phenotype was reversed as inflammation worsened (DSS-5 and DSS-7) (Fig.2d). Immunoblots of whole tissue lysates isolated from control and DSS-7 mice revealed significant NCoR1 upregulation in DSS-7 compared to control mice (Fig S2c). Notably, however, ∼ 4-fold downregulation of NCoR1 expression was seen at the transcriptional level (Fig S2d). Crypts were isolated from the colon of various mice groups and subjected to immunoblotting. LGR5 protein, a marker of crypt resident stem cells^17^ was present only in isolated crypt, but not in whole tissue lysate (data not shown). In isolated crypts, NCoR1 was significantly downregulated in DSS-3 and DSS-5 grouped mice compared to control groups (Fig.2e). This indicated a lowering of NCoR1 in the colon crypts during the onset of inflammation. Immunohistochemistry also indicated inflammation to be progressively higher in tissue sections of DSS-3 versus DSS-5 versus DSS-7 as evident by morphological and histopathological markers including loss of GC, immune cell infiltration, crypt loss, which were absent in control. Notably, NCoR1 presence was observed in the colon crypts in the nuclear region of control, DSS3 and DSS-5 but not DSS-7. However, in DSS-7, NCoR1 presence was observed only in infiltrating immune cells (Fig.2f).

**Fig 2:**
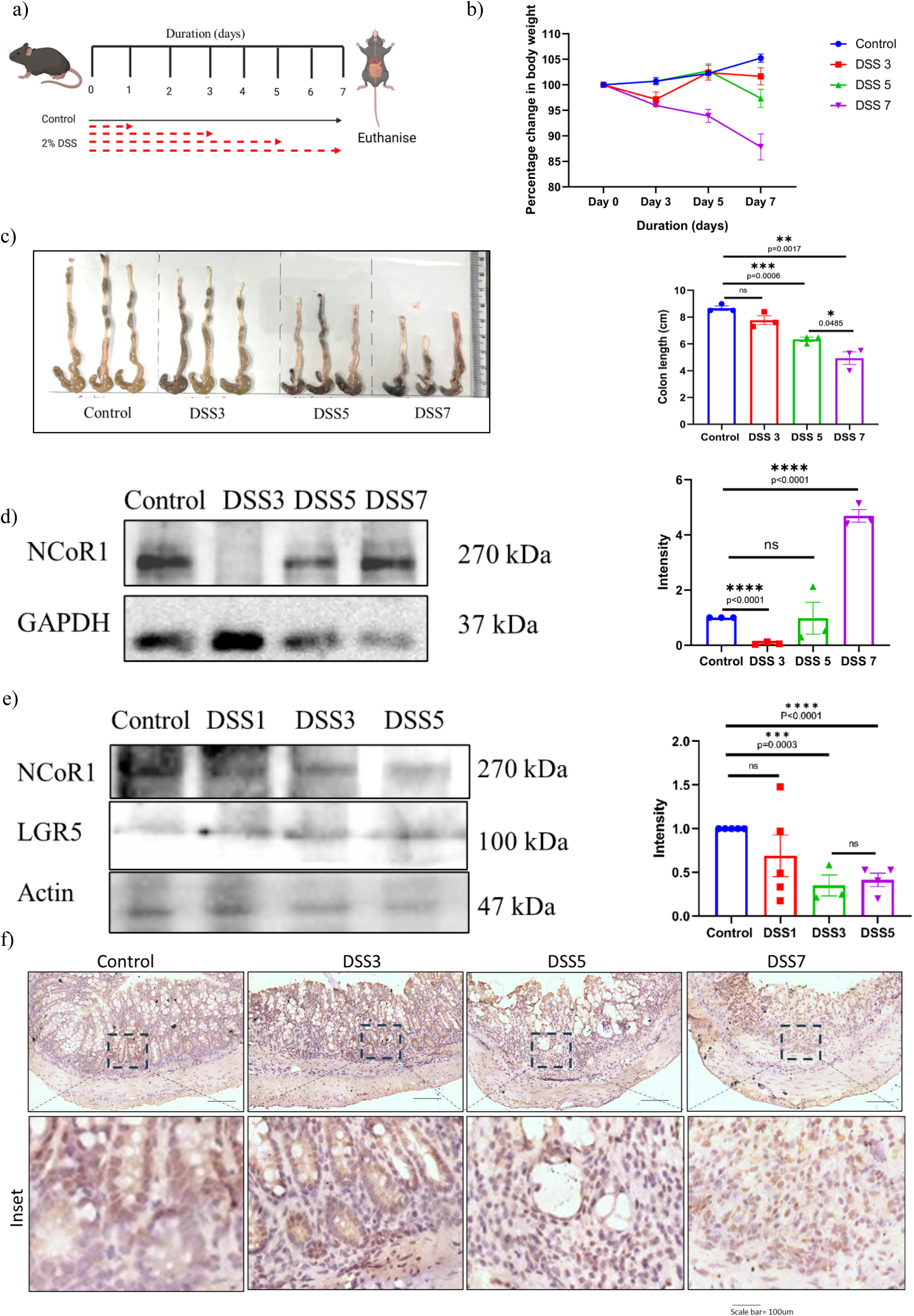
Expression dynamics of NCoR1 in murine colitis. a) Schematic representation of the experimental plan for DSS dynamics in C57Bl/6 mice. Point of DSS administration indicated by red arrows (Created with BioRender.com). b) Graph shows percent body weight change in Control, DSS3, DSS5 and DSS7mice groups. c) Gross morphology of colon and caeca of respective mice groups. The graph on the right shows colon length quantification. d) Immunoblot of NCoR1 protein in whole tissue lysate of DSS dynamics Control, DSS3, DSS5 and DSS7 mice. The graph on the right represents densitometric analysis showing fold intensity of NCoR1 expression calculated by normalising to loading control (GAPDH). e) Immunoblot of NCoR1 protein in crypt lysate Control, DSS1, DSS3 and DSS5. The graph on the right represents densitometric analysis showing fold intensity of NCoR1 expression calculated by normalising to loading control (Actin). f) Representative immunohistochemistry images of colon from control, DSS3,DSS5 and DSS7 mice stained for NCoR1 (n=3 per group) using anti-NCoR1 antibody. (scale bar=100um) .Zoom in images are shown in the inset. Each dot represents (c, d, e) one mouse. All data were expressed as means ± SEM, unpaired Student t-test was used to calculate statistical significance \**P* < 0.05; \*\**P* < 0.01; \*\*\**P* < 0.001; \*\*\*\**P* < 0.0001; ns, not significant. See also Fig S2, Fig S3

These results were revalidated in IL10^-/-^ mice, which tend to develop spontaneous inflammation, thereby serve as a genetic model for IBD^18^. In order to obtain a synchronised inflammation, these mice were subjected to a mild inflammatory trigger (0.5% DSS in drinking water) (Fig.S3a,b). IL10^-/-^ mice no significant change in the colon was observed post euthanasia (Fig.S3c), signs of very mild inflammation were evident in the colon sections (Fig.S3e). Immunoblots of whole tissue revealed significant downregulation of NCoR1 in the IL10^-/-^ mice compared to control mice (Fig.S3d). These were similar to the DSS-3 mice with mild inflammation described in Figure 2. Altogether, these data suggest NCoR1 expression is dynamically regulated; it undergoes downregulation in intestinal crypts during colitis, suggesting a role in gut inflammation.

### NCoR1 modulates GC function by suppressing MUC2 expression

Intestinal crypts harbour several types of cells, all of which are produced by the same pool of resident stem cells. These cell types include transit amplifying cells, goblet cells, enterocytes etc^19^. We set forth to investigate the possible role of NCoR1 in the differentiation of GC colonic HT-29 cell culture model. As reported earlier^4^, post-treatment with 250 mM galactose-containing DMEM media for 3 days (goblet cell differentiation media) and allow them to differentiate into goblet-like cells (Fig.3a). These cells display goblet cell-like features, such as presence of vacuolated morphology (Fig.3b), upregulation of GC markers including MUC2 (∼100-fold upregulated), MUC5AC and downregulation of stem cell marker LGR5 (∼5-fold downregulation) (Fig.S4a). These cells therefore were designated as HT29^Diff^, while the parent line were called as HT29^Undiff^. Immunoblots showed significant upregulation of NCoR1 and GC marker- CLCA1 in HT29^Diff^ cells compared to HT29^Undiff^ (Fig.3d). Next, stable NCoR1 knock-down were created in HT29 cells (HT29^Undiff-kd^ and HT29^Undiff-C^), Morphological changes along with presence of multiple vacuoles observed in case of HT29^Diff-C^ when subjected to GC differentiation media. Notably, these features were significantly reduced in HT29^Diff-kd^ despite treatment with differentiation media (Fig.3e), thereby indicating a role for NCoR1 in GC differentiation. Furthermore, MUC2 expression was upregulated upon treatment in HT29^Diff-C^ when subjected to differentiation media, but not in case of HT29^Diff-kd^ (Fig.3f). Similarly, MUC2 was also downregulated in the NCoR1 stably knockdown in GC line HT29 MTX (HT29 MTX^NCoR1kd^) when compared to control (HT29 MTX^scr^) (Fig.S4b) Presence of NCoR1expression was also seen in other GC lines, LS174, HT29^Diff^, and HT29 MTX cells, but not in non-goblet cell (epithelial cell lines) HCT8 and HT29^Undiff^ (Fig.S4c). Macrophage line J774 included as a non-relevant control also showed NCoR1 expression. Taken together, NCoR1 modulates GC differentiation and function. Presence of NCoR1 is required for GC maturation and MUC2 production.

**Fig. 3:**
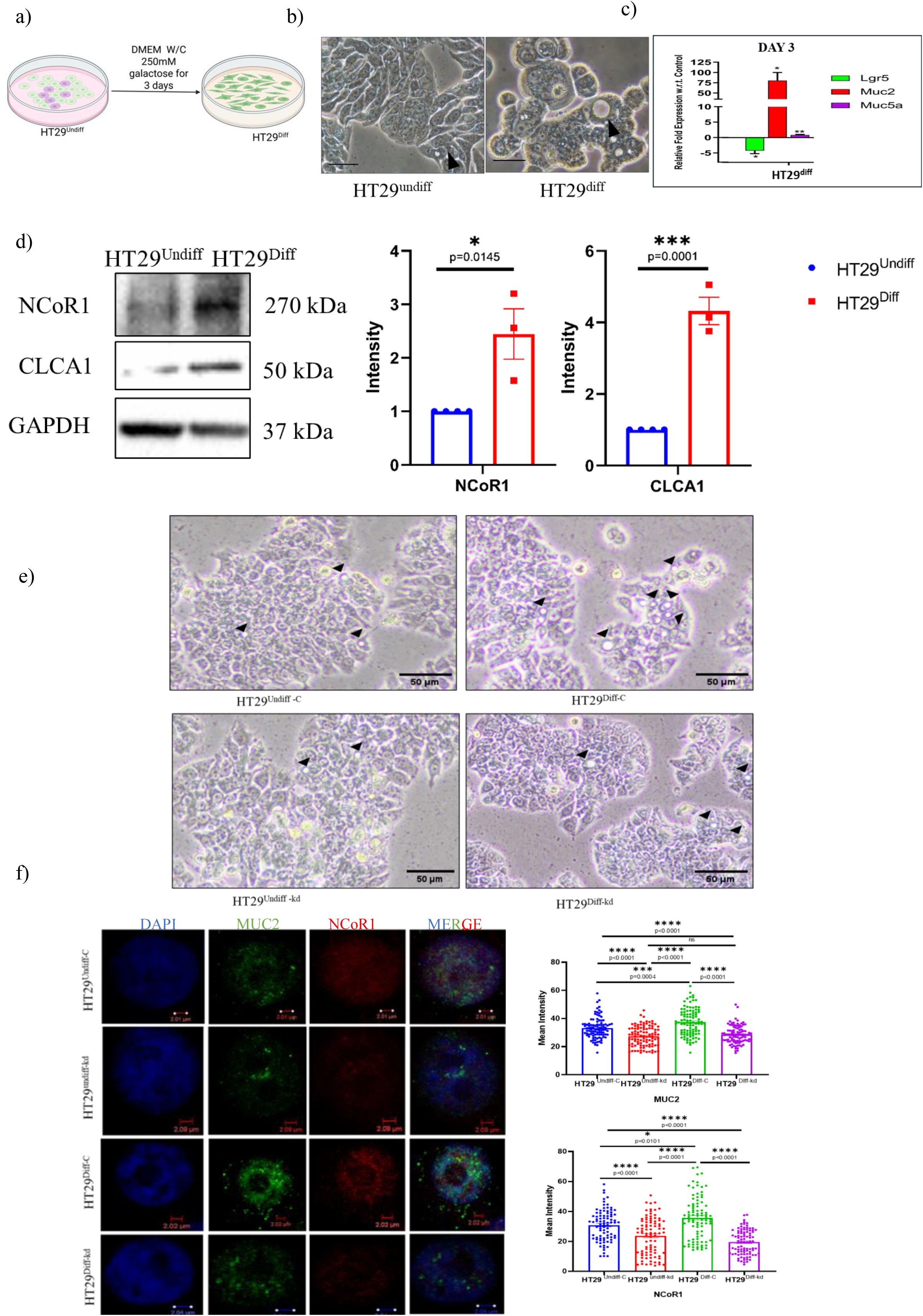
Regulation of goblet cells by NCoR1. a) Schematic representation of the experimental plan for galactose-mediated HT29 pluripotent cells into goblet cells differentiation.(Created with BioRender.com). b) Brightfield images showing goblet cell morphology in HT29^Undiff^ and HT29^Diff^ cells (Scale bar =50um). c) qRT -PCR analysis of relative fold expression of LGR5, MUC2 and MUC5AC genes in HT29^Diff^ cells with respect to HT29^undiff^ cells as control. HPRT was used for normalisation. Experiment performed in triplicate. d) Immunoblotting of NCoR1 and CLCA1 in HT29^Undiff^ and HT29^Diff^ cells. The graph on right represents densitometric analysis showing fold intensity of NCoR1 and CLCA1 expression calculated by normalising to loading control (GAPDH) e) Brightfield images showing goblet cell morphology in HT29^undiff-C^, HT29^Undiff-kd^, HT29^Diff-C^ and HT29^Diff-kd^ cells. Superscript C represents nontargeted knockdown control and superscript kd represents NCoR1 stable knockdown. Scale bar =50um. f) Representative Confocal images of NCoR1 (red) using anti-NCoR1 antibody and MUC2 (green) using anti-MUC2 antibody in HT29^undiff-C^, HT29^Undiff-kd^, HT29^Diff-C^ and HT29 ^Diff-kd^ cells (n=90) (scale bar=2 um). Graphs on the right shows NCoR1 and MUC2 fluorescence intensity measured. Each dot represents (d) individual experiment. All data were expressed as means ± SEM, unpaired Student t-test was used to calculate statistical significance \**P* < 0.05; \*\**P* < 0.01; \*\*\**P* < 0.001; \*\*\*\**P* < 0.0001; ns, not significant. See also Fig S4

### Downregulation of NCoR1 aggravates intestinal inflammation in murine colitis

NCoR1 global knockout mice are nonviable^20^, therefore an *in vivo* knockdown model of NCoR1 was utilised to study its role in gut inflammation. A customised, morpholino based NCoR1 specific knockdown model was developed (Gene Tools LLC, USA). A non-targeting morpholino against human beta-globin, that is absent in mice, was also included as a negative control. Mice were divided into four different groups: control (Ctrl), DSS, NCoR1 knockdown (Ctrl^NCoR1KD^) and NCoR1 knockdown+ DSS (DSS^NCoR1KD^) (Fig.4a). Treatment of the morpholino Ctrl and Ctrl^NCoR1KD^ did not result in weight loss or any other discernible change (Fig 4b). After the stipulated course, the animals were euthanized and various tissues of interest were harvested. Notably, a significant reduction in the colon length in DSS and DSS^NCoR1KD^ was observed, while those from Ctrl and Ctrl^NCoR1KD^ groups appeared normal (Fig 4c). Splenomegaly was also evident only in DSS and DSS^NCoR1KD^ groups (Fig 4d). qRT-PCR analysis revealed a ∼2-fold downregulation of NCoR1 in Ctrl^NCoR1KD^ and DSS, and ∼10-fold downregulated in DSS^NCoR1KD^ with respect to Ctrl, confirming successful knockdown of NCoR1 in the mice. Expression of MUC2 was ∼2-fold downregulated in DSS, ∼2-fold downregulated in Ctrl^NCoR1KD^ and ∼3-fold downregulated in DSS^NCoR1KD^ mice in comparison to Ctrl (Fig.4e). Downregulation of NCoR1 was also seen in the spleen (Fig.4f), indicating a global knockdown of NCoR1. Notably, the extent of inflammation was more severe in the DSS^NCoR1KD^ mice when compared to DSS (Fig.4g). Number of GCs per crypt, based on Alcian blue staining pattern, revealed reduction in the Ctrl^NCoR1KD^ (∼10cells/crypt), and severely reduced in DSS (∼ 5cells/crypt) and DSS^NCoR1KD^ (∼2.5cells/crypt) when compared to Ctrl (∼15cells/crypt) (Fig.4h). Taken together the data shows that deficiency of NCoR1 leads to an exacerbated inflammation accompanied with marked reduction of GCs in NCoR1 knockdown model of colitis.

**Fig 4:**
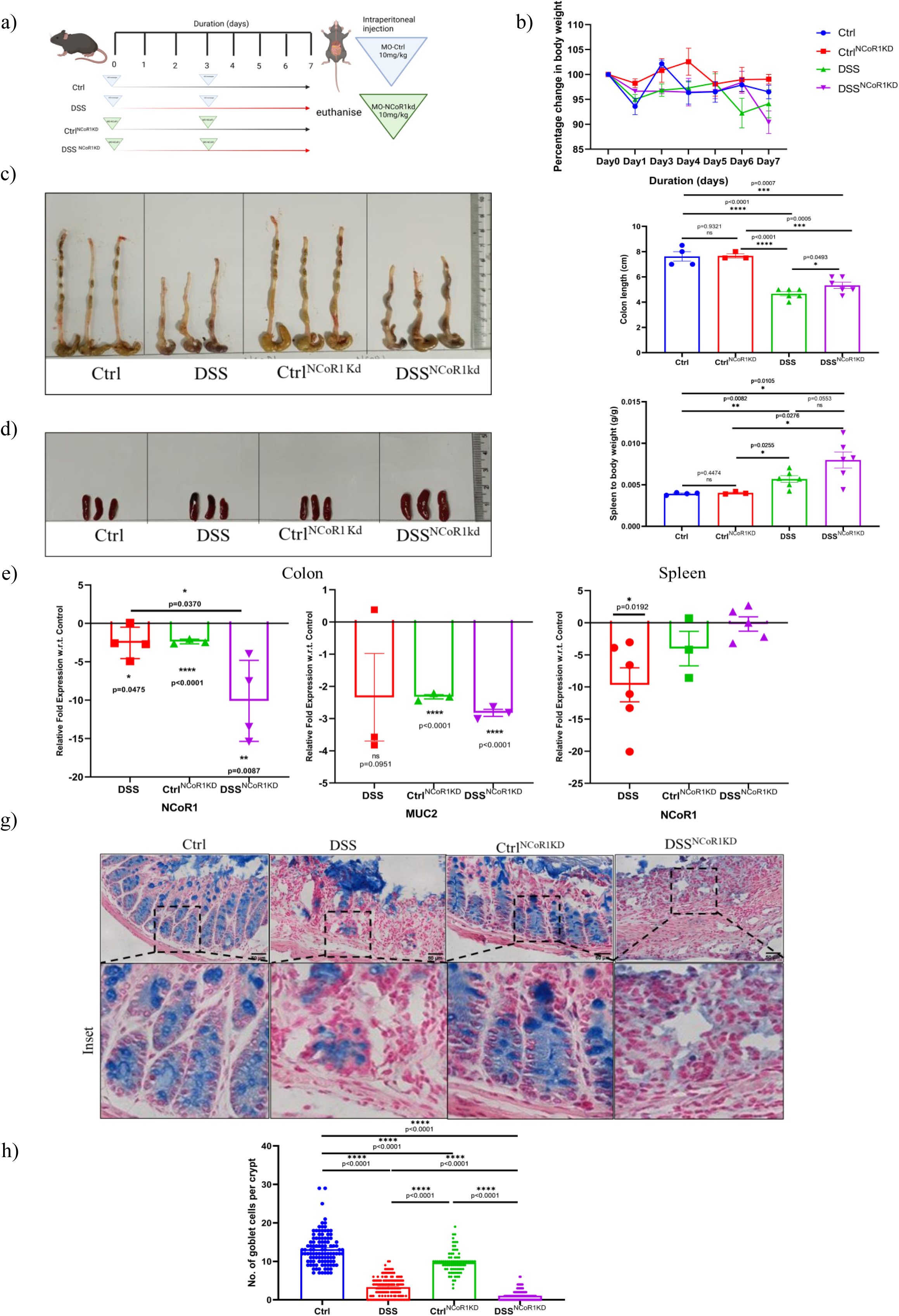
Downregulation of NCoR1 aggravates intestinal inflammation in murine colitis. a) Schematic representation of the experimental plan for knockdown of NCoR1 in C57Bl/6 mice using vivo-morpholino w/c or w/o 2% DSS in drinking water. Point of administration of DSS (Red) control morpholino (blue) and NCoR1 Morpholino(green) is indicated by respective arrows. Morpholino(s) were administered via intraperitoneal injection on the mentioned point of administration (Created with BioRender.com). b) Graph shows percentage body weight change in the mice groups Ctrl, DSS, Ctrl^NCoR1kd^ and DSS^NCoR1kd^. c) Gross morphology of colon and caeca of indicated mice group. Graph on the right shows colon length quantification. d) Gross morphology of spleen of indicated mice group. Graph on the right shows spleen to body weight ratio. e) qRT-PCR analysis of relative fold expression of NCoR1 and MUC2 genes in DSS, Ctrl^NCoR1kd^ and DSS^NCoR1kd^ relative to average control values of Ctrl in colon tissues. GAPDH was used for normalisation. f) qRT-PCR analysis of relative fold expression of NCoR1 gene in DSS, Ctrl^NCoR1kd^ and DSS^NCoR1kd^ relative to average values of Ctrl in spleen tissues. GAPDH was used for normalisation g) Representative alcian blue staining in colon tissue sections of Ctrl, DSS, Ctrl^NCoR1kd^ and DSS^NCoR1kd^ mice (n=4 per group) (scale bar=50um). Zoom in images are shown in the inset of the indicated area. h) graph showing goblet cell count in the crypts of Ctrl, DSS, Ctrl^NCoR1kd^ and DSS^NCoR1kd^ mice colon (n=120). Each dot represents (c,d,e,f) one mouse. All data were expressed as means ± SEM, unpaired Student t-test was used to calculate statistical significance \**P* < 0.05; \*\**P* < 0.01; \*\*\**P* < 0.001; \*\*\*\**P* < 0.0001; ns, not significant.

### Induction of NCoR1 via Vitamin D3 increases GC numbers and alleviates murine colitis

Vitamin D is commonly administered to UC patients as an adjunctive therapy^21,22^ while NCoR1 is known to interact with the Vitamin D Receptor (VDR) to mediate gene silencing. Both of these proteins are known to regulate each other in different systems^23,24^, but their role is not explored in the context of goblet cells. HT29-MTX cells subjected to 25 μM vitamin D3 showed significant upregulation of NCoR1 (Fig.S5a). To further investigate this, a 3-week vitamin D3-infused diet model in mice was established. Mice were categorised into four groups i.e. control group (Ctrl), vitamin D3 group (Ctrl^VD^) (fed with 3000IU/kg with 1% CaCl_2_ in chow for 3 weeks), DSS (2% DSS in the last 5 days of 3^rd^ week), vitamin D3+ 2.5% DSS (DSS^VD^) (mice fed with Vitamin regimen as stated above along with 2% DSS in the last 5 days) (Fig.5a). Post treatment, no weight loss or discernible change was observed in Ctrl^VD^ and Ctrl. Mice were euthanised post 3 weeks from the start of experiment (Fig 5b). Notably, there was a significant reduction in the colon length in the DSS mice compared to Ctrl, Ctrl^VD^ or DSS^VD^ mice (Fig.5c). Splenomegaly was observed in DSS and DSS ^VD^ groups compared control (Fig.S5c). While it was non-significant, the extent of splenomegaly was slightly less in DSS ^VD^ compared to DSS. Immunoblots of the whole tissue lysates depicted a significant increase in NCoR1 in the DSS group with respect to Ctrl, though a non-significant increase was seen in the Ctrl ^VD^ and DSS^VD^ (Fig.5d). NCoR1 levels were maintained in the crypts of Ctrl^VD^ and Ctrl groups, downregulated in the DSS^VD^ in comparison to control groups, but was significantly upregulated when compared to DSS. Enriched levels of LGR5 confirmed successful crypt isolation (Fig.5e). Immunohistochemistry (IHC) Alcian blue – Nuclear fast red staining showed discernible signs of inflammation in DSS and DSS^VD^ mice by morphological and histopathological markers compared to Ctrl, Ctrl^VD^ (Fig.5f). Signals of NCoR1 expression was seen in crypts of Ctrl and Ctrl^VD^. Interestingly, expression of NCoR1 was seen to be higher in crypts of Ctrl^VD^ compared to Ctrl mice. In DSS higher expression of NCoR1 was seen in the infiltrated immune cells. However, in DSS^VD^ crypts were rescued, and expression of NCoR1 was seen in the crypts. Furthermore, GC count per ∼120 crypts in each per was estimated. The number of GCs were significantly reduced in DSS mice (∼2 cells/crypt) with respect to controls (∼10 cells/crypt), while it increased in Ctrl^VD^ mice. In DSS^VD^ mice (∼5 cells/crypt), the numbers were significantly higher than the number of goblet cells present per crypt of DSS mice (Fig.5g). This suggests that Vitamin D3 maintains NCoR1 levels in the crypts of the colon thereby protecting goblet cell numbers and rescues colitis.

**Fig 5:**
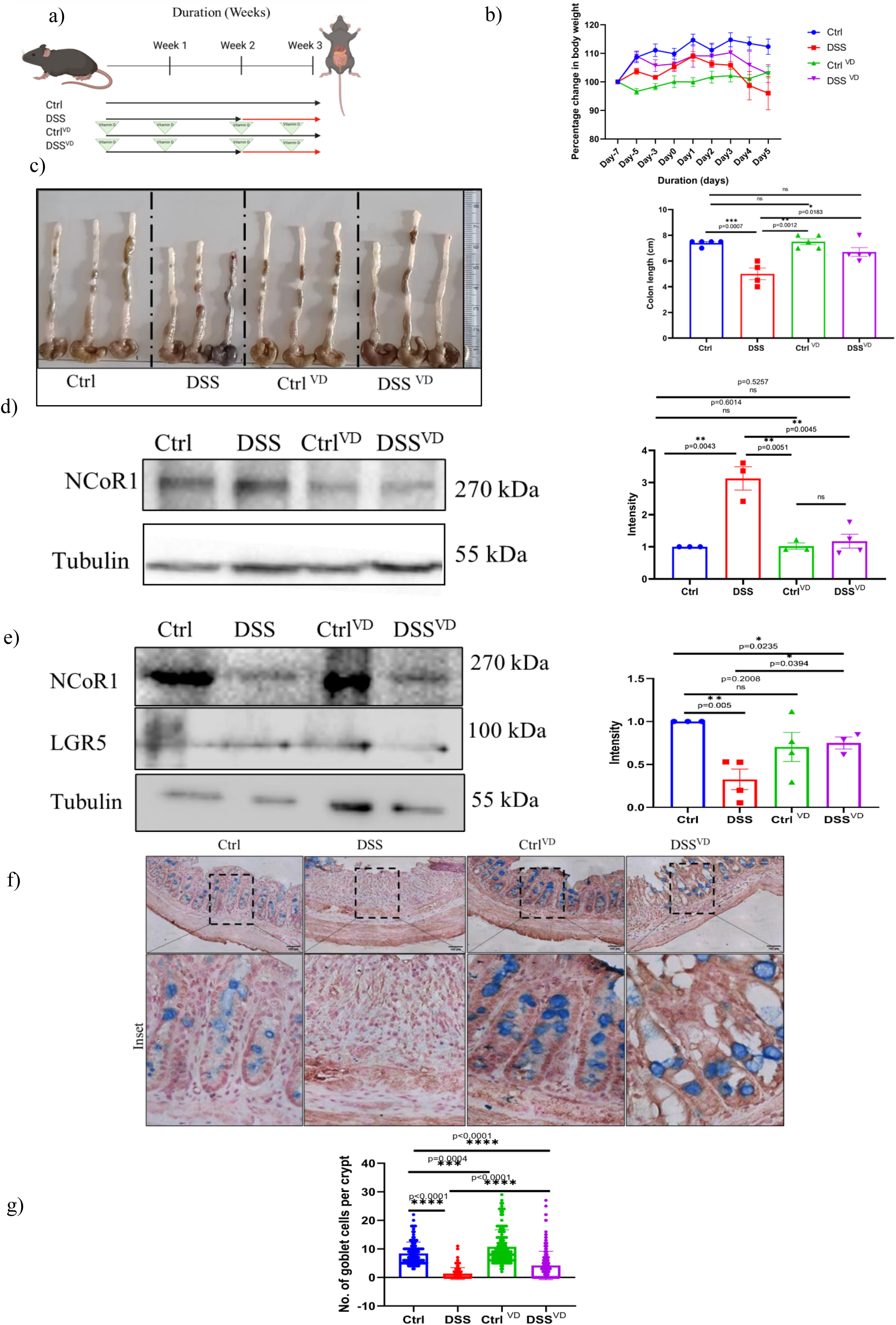
Upregulation of NCoR1 via Vitamin D3 in c57BL/6 mice. a) Schematic representation of the experimental plan for vitamin D-mediated overexpression of NCoR1 in C57Bl/6 mice w/c or w/o 2% DSS in drinking water. Vitamin D was mixed in feed at a dosage of 3000 IU/kg/day with 1% CaCl_2_ throughout experiment. Duration of administration of DSS (red), and vitamin D (green triangle) are indicated by respective arrows (Created with BioRender.com) b) Graph showing the percentage body weight change in Ctrl, DSS, Ctrl^VD^ and DSS^VD^ mice groups. c) Gross morphology of colon and caeca of Ctrl, DSS, Ctrl^VD^ and DSS^VD^ mice. Graph on the right showing colon length quantification. d) Immunoblot of NCoR1 protein in whole tissue lysate of Ctrl, DSS, Ctrl^VD^ and DSS^VD^ mice. The graph on the right represents densitometric analysis showing fold intensity of NCoR1 expression calculated by normalising to loading control (Tubulin) e) Immunoblot of NCoR1 protein in crypt lysate of Ctrl, DSS, Ctrl^VD^ and DSS^VD^ mice. The graph on the right represents densitometric analysis showing fold intensity of NCoR1 expression calculated by normalising to loading control (Tubulin). f) Representative immunohistochemistry (IHC-Alcian blue) images of colon from Ctrl, DSS, Ctrl ^VD^ and DSS ^VD^ mice stained for NCoR1 (n=4 per group) (scale bar=100um). g) Graph showing goblet cell count in the crypts of Ctrl, DSS, Ctrl^VD^ and DSS^VD^ mice (n=120 crypts per group). Each dot represents (c,d,e) one mouse. All data were expressed as means ± SEM, unpaired Student t-test was used to calculate statistical significance \**P* < 0.05; \*\**P* < 0.01; \*\*\**P* < 0.001; \*\*\*\**P* < 0.0001; ns, not significant. See also fig S5a and fig S5c.

### KLF16 acts as a potential target under the regulation of NCoR1

NCoR1 engages with HDACs to control gene regulation and govern multiple cellular process^25,26^. However limited information is available regarding its role in intestinal goblet cells. To understand this regulation, HT29-MTX cells were subjected to 5 μM Tricostatin A (TSA: pan HDAC inhibitor^27,26^) for 24 hrs followed by immunoblotting of lysates. TSA treatment led to significant downregulation of NCoR1 (Fig.S5b). This suggests that expression of NCoR1 can be altered upon targeting HDACs. TSA-treated HT29-MTX cells were subjected to Chromatin Immunoprecipitation followed by sequencing (ChIP-Seq). Chromatin were pulled down using NCoR1 antibody from the control, and TSA-treated HT29-MTX cells. The genes that exhibited differential expression in control and TSA-treated HT29-MTX cells were visualized in respective volcano plots. Genes that showed significant differential expression were identified as significantly differentially bound (FDR <=0.05) and were marked in pink. (Fig.6a), Heat maps were generated to identify genes that are specifically bound by NCoR1 (Fig.6b). Four targets, AGAP3, KLF16, EXOSC6 and GLTPD2, were chosen based on fold change and their potential to regulate different aspects of goblet cells (TABLE 2). The role of any of these molecules is not known with regard to the GC. ChIP-qRT-PCR was performed against these targets in previously mentioned HT29^Undiff^ and HT29^Diff^ cells. Antibody-dependent chromatin were pulled down using NCoR1 antibody, along with IgG as a negative control. AGAP3 and GLTPD2 showed higher binding in HT29^Undiff^ NCoR1 in comparison to HT29^Diff^ NCoR1, showing the potential to regulate the undifferentiated cells; EXOSC6 showed almost no change in any condition. But, KLF16 showed increased binding in the HT29^Diff^ NCoR1 in comparison to HT29^Undiff^ NCoR1, showing the potential to regulate the differentiated goblet cells (Fig S6a). KLF16 was further validated by ChIP qRT-PCR in HT29-MTX control and TSA-treated cells. Upon downregulation of NCoR1 by TSA, a concomitant reduction in the binding of NCoR1 to KLF16 was observed. Whereas there was no change in the IgG and histone H3 (H3) controls (Fig.6c), as also depicted in the heat map.

**Table 1:**
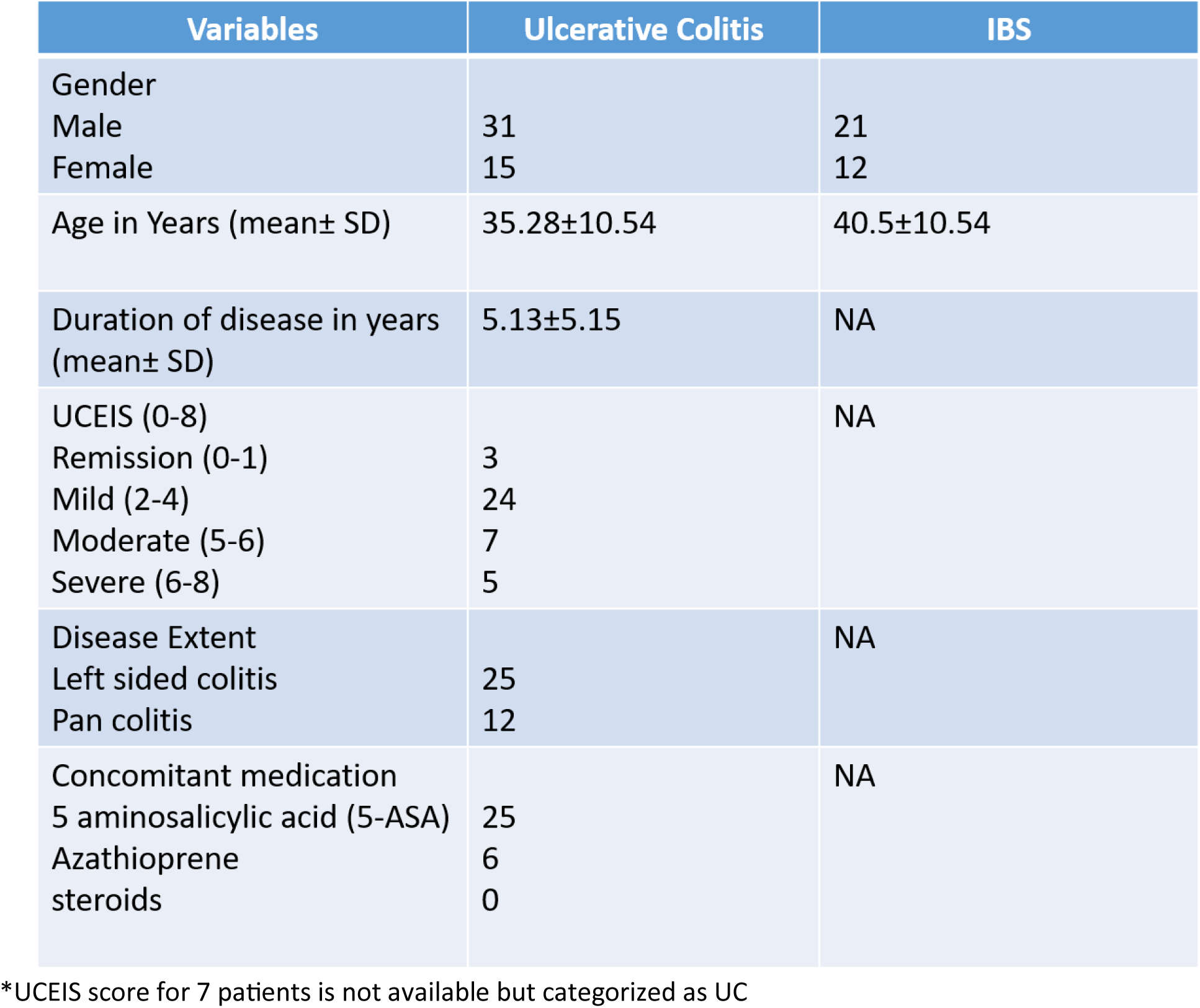
Patient details: Details of the patient’s clinical parameters.

**Table 2:** List of potential interactors of NCoR1.

| Gene Name | Gene Function | Reference |
| --- | --- | --- |
| AGAP3 | GTPase-activating protein | <sup>51</sup> |
| KLF16 | Transcription factor | <sup>42</sup> |
| EXOSC6 | RNA regulation | <sup>52</sup> |
| GLTPD2 | Glycolipid transfer protein | <sup>53</sup> |

**Table 3:** List of Antibodies.

| S.No. | Item | Catalogue | Make |
| --- | --- | --- | --- |
| 1. | Rabbit anti-NCoR1 Antibody | 5948 | Cell Signalling Technologies |
| 2. | Mouse anti-KLF16 Antibody (F-4) | sc-377519 | Santa Cruz Biotechnology |
| 3. | Mouse anti-MUC2 | MA5-12345 | Invitrogen |
| 4. | Rabbit anti- NCoR1 | PA1-844A | Invitrogen |
| 5. | Mouse anti-GAPDH antibody | 39-8600 | Invitrogen |
| 6. | Rabbit anti-β-Actin antibody | 4970S | CST |
| 7. | Goat anti-Rabbit-IgG (HRP) antibody | 111-035-008 | Jackson Laboratories |
| 8. | Goat anti-mouse-HRP conjugated antibody | 115-035-008 | Jackson Laboratories |
| 9. | Goat anti-Rabbit IgG (Alexa Fluor 488) antibody | 211-032-171 | Jackson Laboratories |
| 10. | Goat anti-Mouse IgG (Alexa Fluor 488) antibody | 115-545-003 | Jackson Laboratories |
| 11. | Goat anti-Rabbit IgG (CyTM5) antibody | 111-175-144 | Jackson Laboratories |
| 12. | Goat anti-Mouse IgG (CyTM5) antibody | 115-175-146 | Jackson Laboratories |
| 13. | Anti-Histone H3 (acetyl K9 + K14 + K18 + K23 + K27) antibody | ab47915 | Abcam |
| 14. | Mouse anti-β-Tubulin monoclonal antibody | T7816 | Sigma-Aldrich |
| 15. | Rabbit anti-CLCA1 antibody | ab180851 | Abcam |
| 16. | Rabbit IgG Polyclonal Antibody | 12-370 | Sigma-Aldrich |
| 17. | mTOR | 2983S | CST |
| 18. | p-mTOR (Ser2448) | 5536S | CST |
| 19. | p-4E-BP1 (Thr37/46) | 2855S | CST |
| 20. | p70S6Kinase | 9202S | CST |
| 21. | p-p70S6Kinase (Thr389) | 97596S | CST |

**Table 4:**
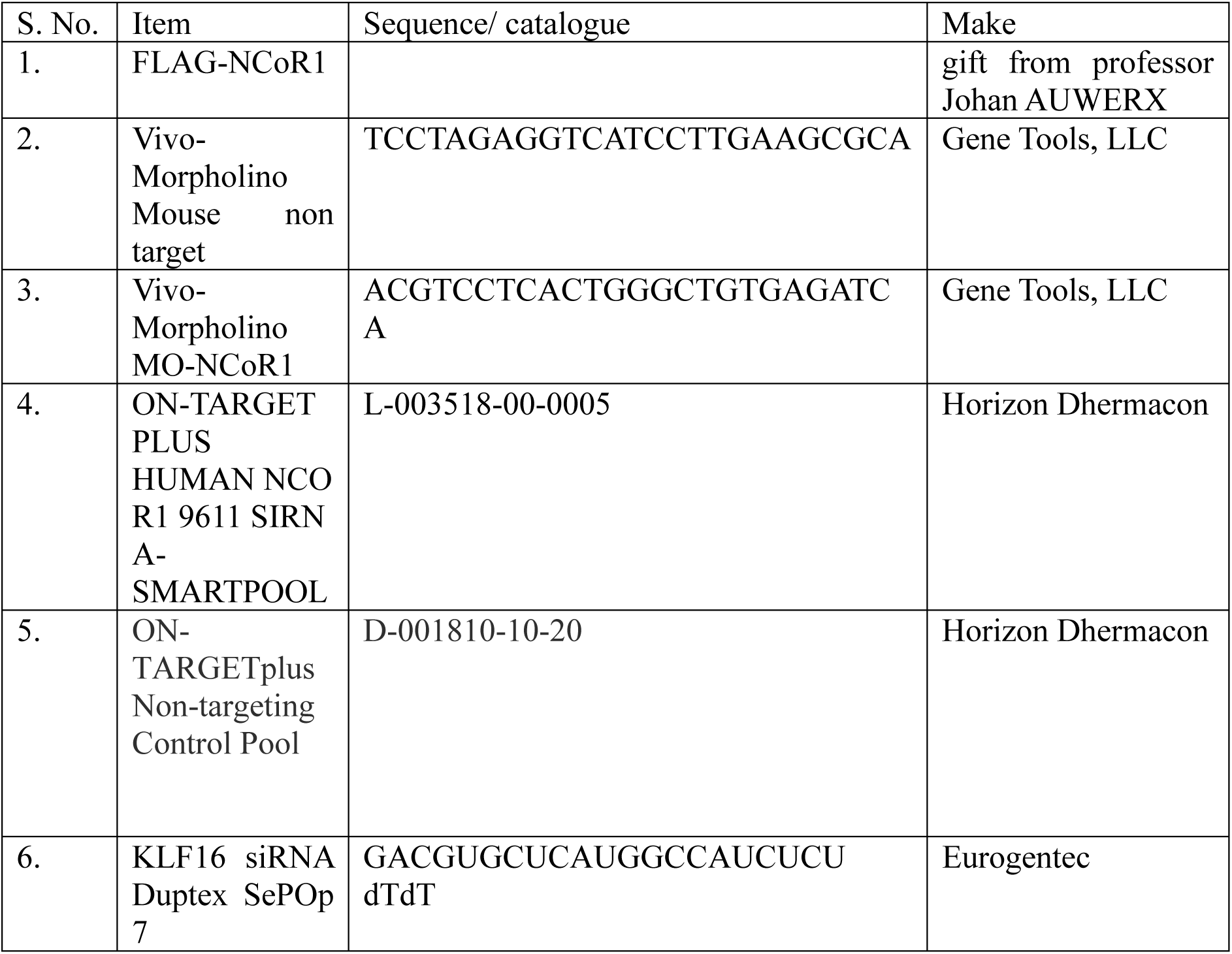
List of Plasmids and siRNA.

**Table 5:**
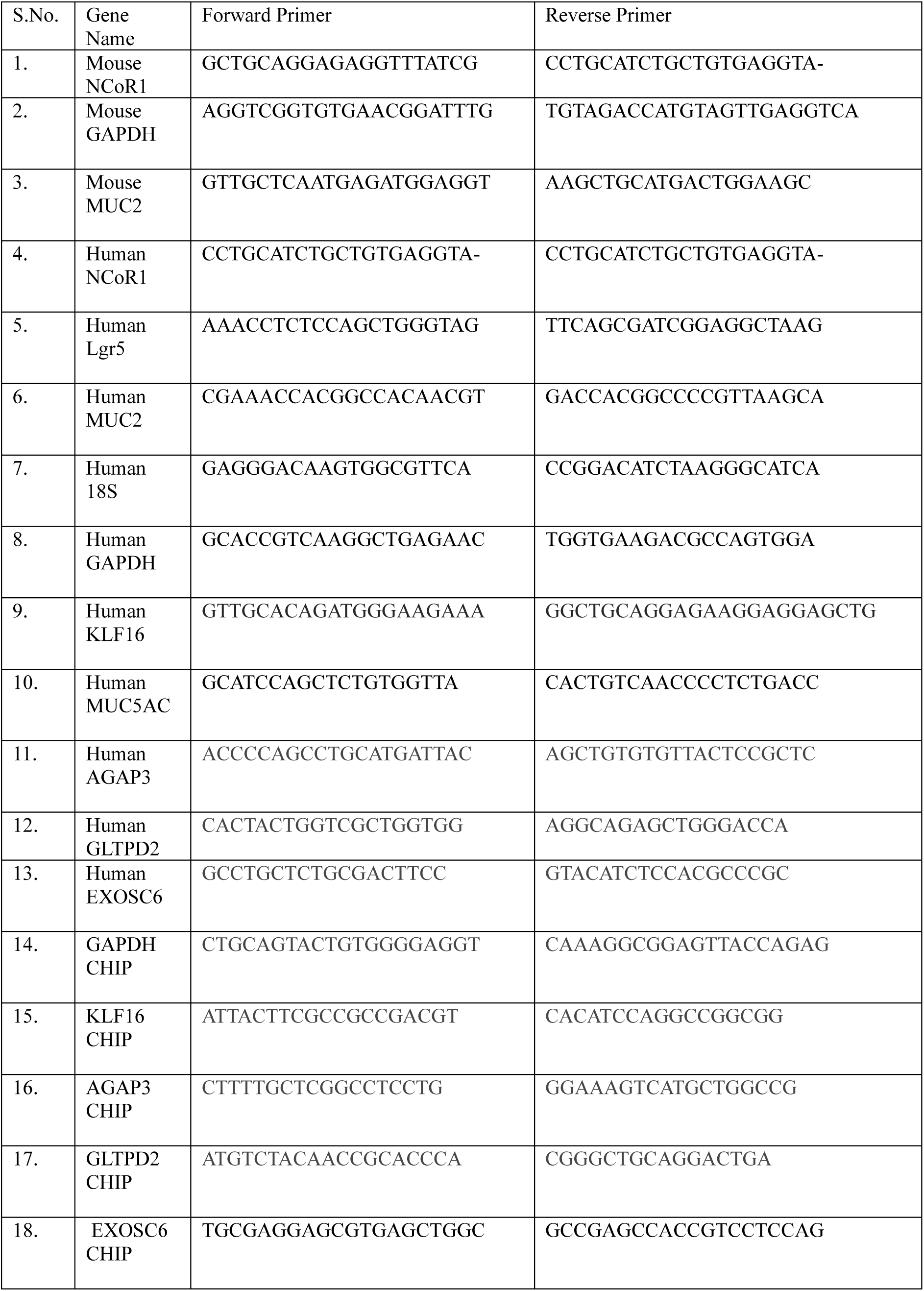
List of Primers.

**Table 6:** List of Reagents.

| S.No. | Item | Catalogue | Make |
| --- | --- | --- | --- |
| 1. | Dextran Sulphate Sodium | 160110 | MP Biomedical |
| 2. | Vitamin D3 | Dr. Exceld3 | Dr. Reddys |
| 3. | SignalStain® DAB Substrate Kit | 8059S | CST |
| 4. | DMEM | 12100046 | Gibco |
| 5. | Fetal Bovine serum (FBS) | 10270-106 | Gibco |
| 6. | Sodium Pyruvate | 11360070 | Gibco |
| 7. | Sodium Bicarbonate | S6014 | Sigma |
| 8. | Penicillin-Streptomycin | 15140122 | Gibco |
| 9. | DMEM no glucose | 11966025 | Gibco |
| 10. | D-(+)-galactose | G0750 | Sigma |
| 11. | DMEM Glutamax | 10566016 | Gibco |
| 12. | TransFectin™ Lipid Reagent | 1703351 | Bio Rad |
| 13. | DHARMAFECT 1 transfection reagent | T-2001-03 | Horizon |
| 14. | OptiMEM | 31985070 | Gibco |
| 15. | CalPhos™ Mammalian Transfection Kit | 631312 | Takara |
| 16. | Tricostatin A | T1952 | Sigma |
| 17. | ProLong™ Diamond Antifade Mountant | P36970 | Invitrogen |
| 18. | RNeasy Mini Kit | 74104 | Quaigen |
| 19. | iScript™ cDNA Synthesis Kit | 1708891 | Bio Rad |
| 20. | iTaq Univer SYBR Green | 1725124 | Bio Rad |
| 21. | RIPA buffer | R0278 | Sigma |
| 22. | Bicinchoninic Acid solution | B9643-1L | Sigma |
| 23. | Nitrocellulose Membrane | 1620115 | Bio Rad |
| 24. | Alcian Blue Solution | B8438 | Sigma |
| 25. | Nuclear Fast Red | GRM2354-1G | HiMedia |
| 26. | Immobilon Forte Western HRP substrate | WBLUF | Merk |

To further investigate the regulation of AGAP3, KLF16, EXOSC6 and GLTPD2, qRT-PCR of these targets were analysed in the control HT29-MTX cells (siCTL) and siRNA-mediated NCoR1 knockdown HT29-MTX cells (NCoR1^KD^). Plasmid-mediated NCoR1 overexpression in HT29-MTX cells (NCoR1^UP^) and the corresponding control cells (NCoR1^CTL^) were also analysed. AGAP3, EXOSC6 and GLTPD2 did not show any significant change in either condition. But KLF16 expression was proportional to NCoR1. As it was significantly upregulated in NCoR1^UP^ and downregulated in NCoR1^KD^ conditions at both transcriptional (Fig.S6b, Fig.S6c) and translational levels (Fig.6d, Fig.6E). To analyse the localisation, both KLF16 and NCoR1, cytoplasmic and nuclear fractions were isolated from the stable NCoR1 knockdown HT29 MTX ^NCoR1kd^ and HT29 MTX^scr^ cells. Both NCoR1 and KLF16 were expressed in the nuclear fraction but not in the cytoplasmic fraction. Downregulation of NCoR1 significantly reduced KLF16 in the nuclear fraction as well (Fig.S6d). Immunofluorescence analysis also revealed the presence of NCoR1 and KLF in the nucleus of the HT29 MTX ^NCoR1kd^ and HT29 MTX^scr^ cells, and downregulation of NCoR1 caused a significant decrease in expression of KLF16 in the nucleus (Fig.6F). Taken together, it was reasonable to conclude that NCoR1 regulates KLF16 in the goblet cells.

**Fig 6:**
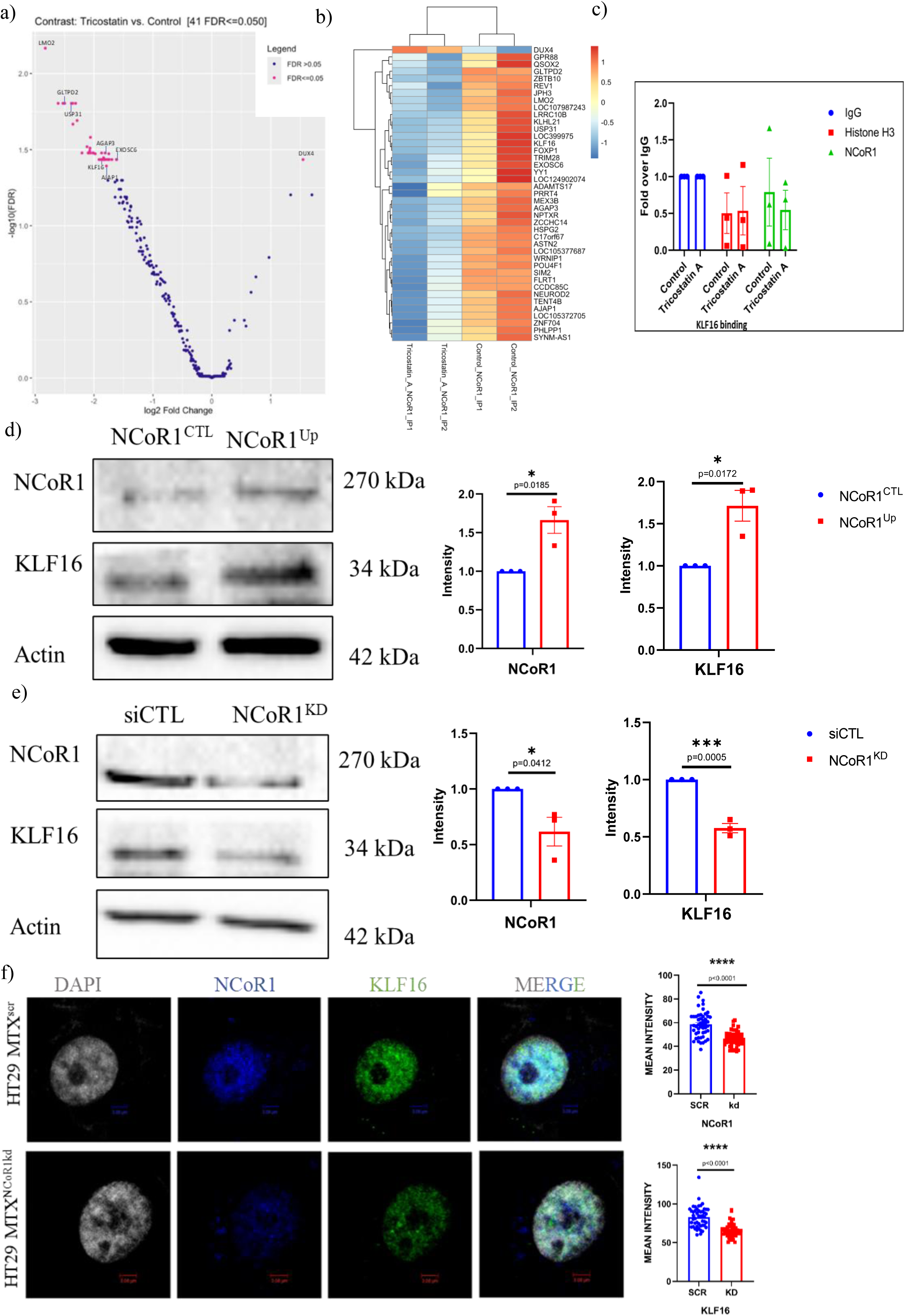
KLF16 is regulated by NCoR1. a) Volcano plot of DEGs representing NCoR1 targets. The pink dots represent genes differentially expressed (adjusted P<0.05) in Tricostatin A-treated and control HT29 MTX cells. The symbol-marked dots indicate the genes with the largest negative or positive standardized mean difference. b) Heat map of genes regulated by NCoR1 in Tricostatin A-treated and control HT29 MTX cells represented in colour contrast. Expression of targets is depicted in the gradient of blue to red, indicating lowest to highest expression, respectively. c) Graph showing chromatin immunoprecipitation of KLF16 binding via chromatin pulled by IgG, H3 and NCoR1 antibodies in Tricostain A-treated and control HT29 MTX cells. Quantification done using the fold-over IgG method. d) Immunoblot of NCoR1 and KLF16 in HT29 MTX cells upon transient overexpression of NCoR1(NCoR1^CTL^ and NCoR1^Up^). The graph on the right represents densitometric analysis showing fold intensity of NCoR1 and KLF16 expression calculated by normalising to loading control (Actin). e) Immunoblot of NCoR1 and KLF16 in HT29 MTX cells upon transient knockdown of NCoR1(siCTL and NCoR1^KD^). The graph on the right represents densitometric analysis showing fold intensity of NCoR1 and KLF16 expression calculated by normalising to loading control (Actin). f) Representative confocal imaging of NCoR1 (blue) stained using anti- NCoR1 antibody and KLF16 (green) stained using anti-KLF16 antibody in HT29 MTX^scr^ and HT29 MTX^NCoR1kd^ cells scale bar=3um. Graphs on the right show fluorescence intensity of NCoR1 and KLF16 measured (n=60). Each dot represents an individual experiment (c, d, and e). All data were expressed as means ± SEM, unpaired Student t-test was used to calculate statistical significance \**P* < 0.05; \*\**P* < 0.01; \*\*\**P* < 0.001; \*\*\*\**P* < 0.0001; ns, not significant.

### KLF16 is required for the expression of MUC2 in GCs

KLF16 is known to regulate endocrine activity at the cellular level to mediate homeostasis. However, the role of KLF16 in colon or goblet cells is not known. To elucidate its role in GC regulation, KLF16 was transiently downregulated in HT29 MTX cells (siCTL and siKLF16). Notably, downregulation of KLF16 decreased NCoR1 at translational (Fig.7a) and transcriptional levels (Fig.7b). Also, expression of MUC2 was analysed by qRT-PCR and immunofluorescence in siCTL and siKLF16 HT29-MTX cells. Downregulation of KLF16 caused ∼3-fold downregulation of MUC2 (Fig.7b), which was also validated by immunofluorescence (Fig.7c). Promoter analysis of MUC2 was done using the pSCAN application, which predicts the binding of different transcription factors to the promoter of MUC2. JASPAR predicted the KLF16 binding site was present at the -88 position of the start site of MUC2 on chromosome 11 (Fig.7d). ∼ -4-fold downregulation of NCoR1 was observed at transcriptional levels upon transient KLF16 knockdown in HT29 MTX cells (Fig.7b), indicating KLF16 regulating NCoR1. Also, JASPAR predicted a KLF16 binding site was present at the -18 position of the promoter on the reverse strand of NCoR1 (Fig.S7). Taken together, NCoR1 and KLF16 regulate each other. Expression pattern of KLF16 in different cell types revealed expression in HT29^Diff^, HT29^Undiff^, HT29 MTX cells and HCT8 but not in macrophage line- J774 (Fig.7e). Taken together, NCoR1 and KLF16 regulate each other. KLF16 binds to the promoter of MUC2, and the downregulation of KLF16 is sufficient to reduce MUC2 expression. This highlights the presence of NCoR1 and KLF16 together, indicating an important axis specified in goblet cell regulation.

**Fig 7.**
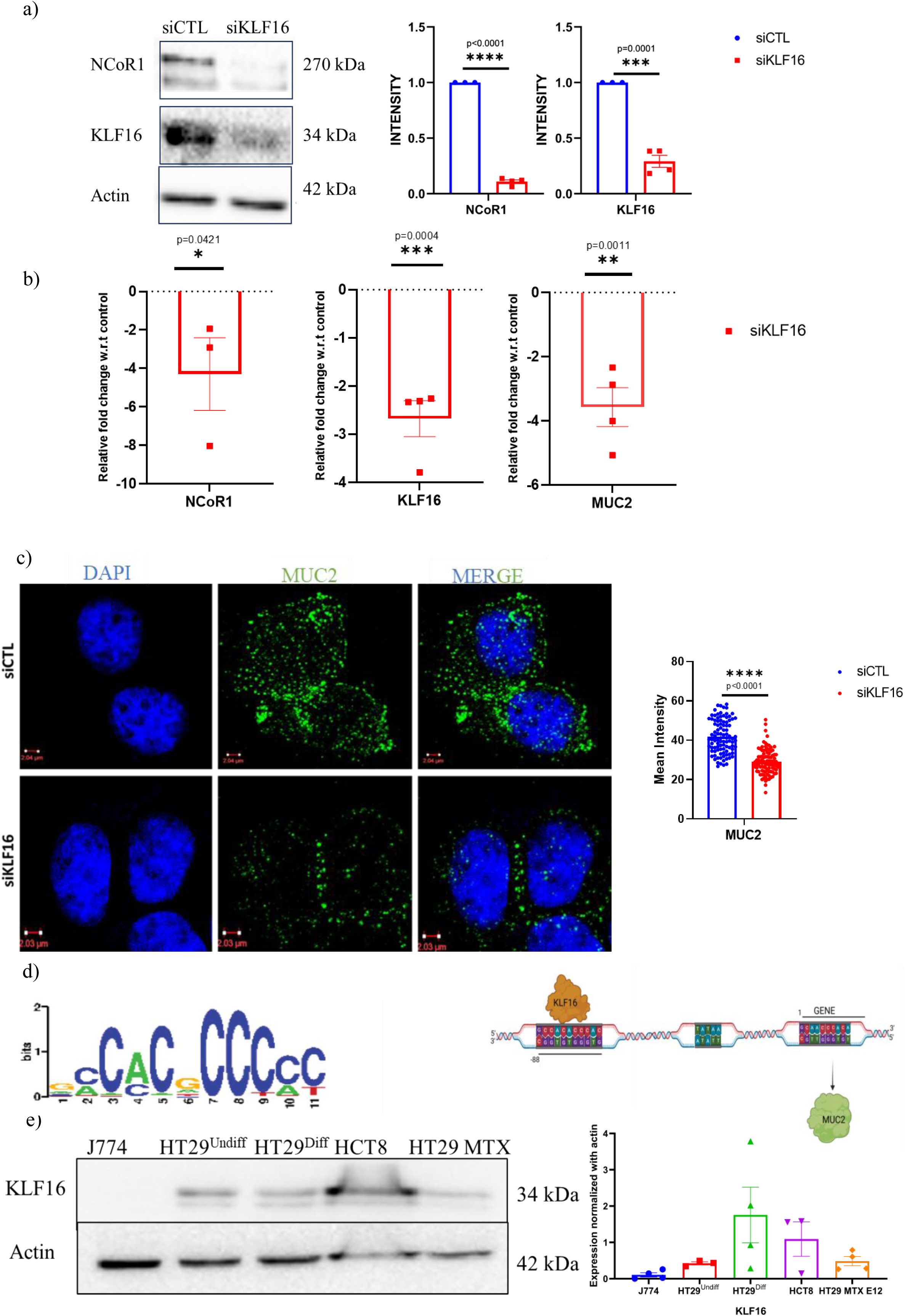
KLF16 regulates NCoR1 and MUC2 in goblet cells. a) Immunoblotting of NCoR1 and KLF16 in HT29 MTX cells upon transient knockdown of KLF16 (siCTL and siKLF16). The graph on the right represents densitometric analysis showing fold intensity of NCoR1 and KLF16 expression calculated by normalising to loading control (Actin). b) qRT-PCR analysis of relative fold expression of NCoR1, KLF16 and MUC2 genes upon transient knockdown of KLF16 (siKLF16) relative to siCTL in HT29 MTX cells. GAPDH was used for normalisation. c) Representative confocal imaging of MUC2 (green) stained using anti-MUC2 antibody in siCTL and siKLF16 in HT29MTX cells (n=90). Scale bar=2 um. The graph shows MUC2 fluorescence intensity measured. d) JASPAR predicted sequence for KLF16 binding. Image on the right indicates pSCAN predicted site for KLF16 binding on MUC2 promoter. Image Created with BioRender.com. e) Immunoblot of KLF16 in J774, HT29^Undiff^,HT29^Diff^, HCT8 and HT29-MTX cell lines. The on the right graph represents NCoR1 expression calculated by normalising to loading control (Actin) specific to each cell type. Each dot represents (a,b and e) individual experiment. All data were expressed as means ± SEM, unpaired Student t-test was used to calculate statistical significance \**P* < 0.05; \*\**P* < 0.01; \*\*\**P* < 0.001; \*\*\*\**P* < 0.0001; ns, not significant.

### Downregulation of expression of KLF16 in murine and human colitis

To examine the expression of KLF16 during inflammation, firstly, protein lysates isolated from the control and UC patients showed ∼-2.5-fold downregulation at transcriptional and translational levels (Fig.8a,b). Similarly, KLF16 displayed a gradual and significant downregulation in the crypts of DSS1, DSS3 and DSS5 in comparison to control (Fig.8c). Immunoblotting of colon tissue lysates from Ctrl, DSS, Ctrl^NCoR1KD^ and DSS ^NCoR1KD^ showed KLF16 was significantly downregulated in Ctrl^NCoR1KD^ and also downregulated in DSS and DSS^NCoR1KD^ with respect to control Ctrl (Fig.8d). Furthermore, KLF16 was downregulated significantly in Ctrl ^VD^, DSS and DSS ^VD^ in comparison to the Ctrl mouse group in the whole tissue (Fig.8e). Crypts lysates also showed significant downregulation of KLF16 in DSS and DSS ^VD^ compared to Ctrl (Fig.8f). Taken together, KLF16 gets downregulated during ulcerative colitis and DSS-mediated colitis. Downregulation of NCoR1 is sufficient to suppress KLF16. Vitamin D-mediated rescue of NCoR1 in the crypts could not rescue KLF16 in the crypt lysates.

**Fig 8.**
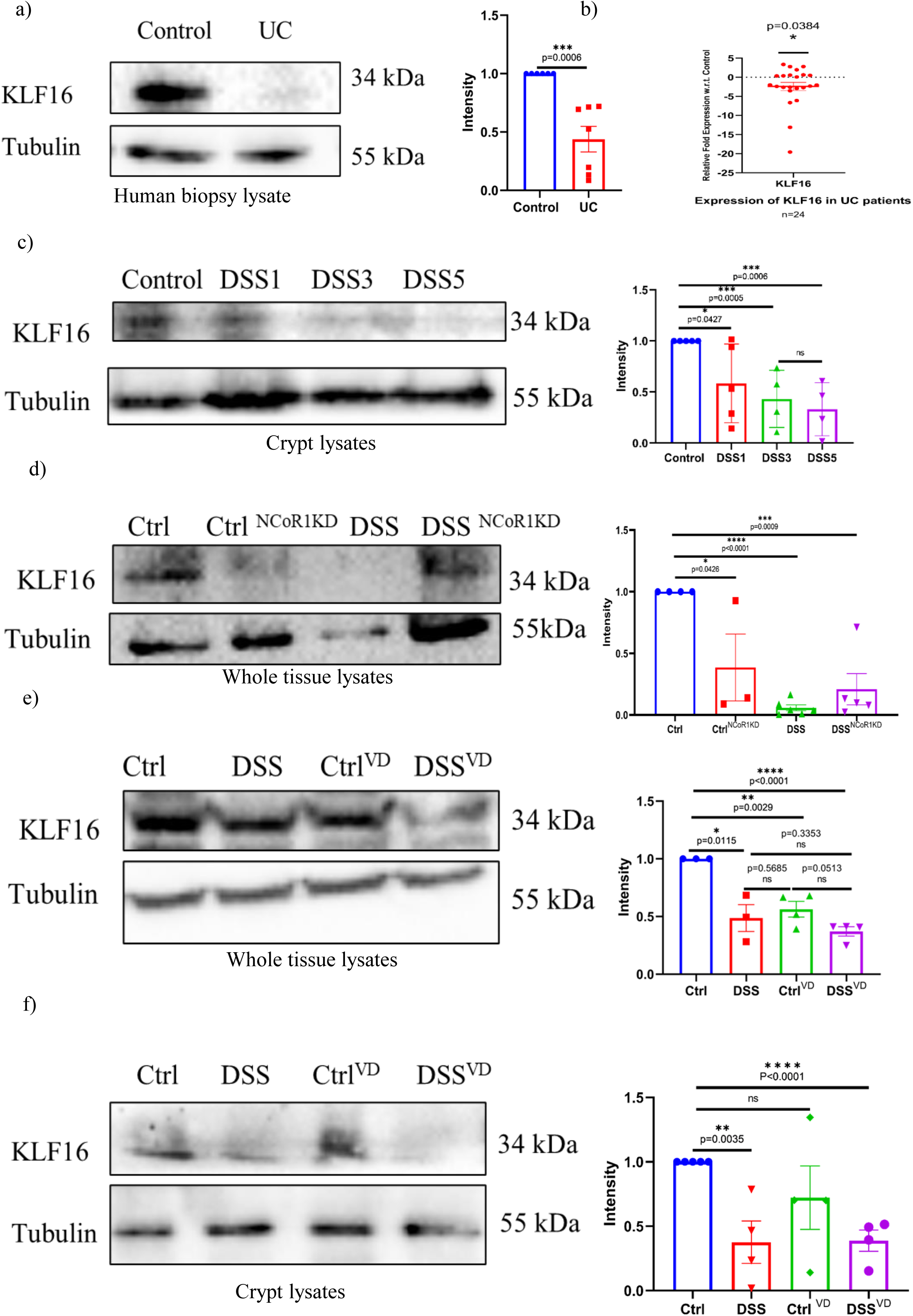
Expression of KLF16 in murine and human colitis. a) Representative Immunoblot of KLF16 protein in human ulcerative colitis (UC) (n = 8) and control (n = 6) biopsy samples. Graph on the right represents densitometric analysis showing fold intensity of KLF16 expression calculated by normalising to loading control (Tubulin). b) qRT-PCR analysis of relative fold expression of KLF16 gene in human UC patient colonic biopsies (n = 24) relative to average control values (n = 23). 18s was used for normalisation. c) Immunoblot of KLF16 protein in crypt lysate of DSS dynamics Control, DSS1, DSS3 and DSS5 mice. The graph on right represents densitometric analysis showing fold intensity of KLF16 expression calculated by normalising to loading control (Actin). d) Immunoblot of KLF16 protein in whole tissue lysate of NCoR1 knockdown mice i.e. Ctrl, Ctrl^NCoR1kd^,DSS and DSS^NCoR1kd^ mice. The graph on right represents densitometric analysis showing fold intensity of KLF16 expression calculated by normalising to loading control (Tubulin). e) Immunoblot of KLF16 protein in whole tissue lysate from the colon of vitamin D-mediated rescue in mice i.e. Ctrl, DSS, Ctrl ^VD^ or DSS ^VD^. The graph on right represents densitometric analysis showing fold intensity of KLF16 expression calculated by normalising to loading control (Tubulin). f) Immunoblot of KLF16 protein in crypt lysate from colon of vitamin D mediated rescue in mice i.e. Ctrl, DSS, Ctrl^VD^ or DSS^VD^. The graph on right represents densitometric analysis showing fold intensity of KLF16 expression calculated by normalising to loading control (Tubulin). Each dot represents (a and b) individual human and (c,d,e and f) individual mice. All data were expressed as means ± SEM, unpaired Student t-test was used to calculate statistical significance \**P* < 0.05; \*\**P* < 0.01; \*\*\**P* < 0.001; \*\*\*\**P* < 0.0001; ns, not significant.

### Downregulation of NCoR1 and KLF16 activates the mTOR pathway and control MUC2 expression in GC

The mTOR1 pathway is known to be upregulated in patients with active UC^28^. Administration of molecules targeting the mTOR pathway acts as a potential treatment for IBD^29^. To check whether the mTOR pathway is activated in the absence of NCoR1, lysates of HT29 MTX^NCoR1kd^and HT29 MTX^scr^ cells were subjected to immunoblotting. KLF16 was significantly downregulated upon the downregulation of NCoR1. There was significant upregulation in p-mTOR, p-pS6, p-4EBP1 and nonsignificant upregulation of mTOR in MTX^NCoR1kd^ with respect to HT29 MTX^scr^ (Fig.9a). Similarly, transient knockdown of KLF16 in goblet cells (siKLF16) downregulated NCoR1. This further activated the mTOR pathway by upregulating p-PS6 but not much change in p-4EBP and mTOR in siKLF16 HT29 MTX with respect to control HT29 MTX cells (Fig.9b). Increased expression of p-mTOR in the NCoR1 knockdown condition indicates activation of mTOR signalling, wherein p-PS6 upregulation in NCoR1 and KLF16 downregulation indicates that the GC maturation is altered upon mTOR activation, leading to defect in mucus secretion and dysbiosis in the intestines. Together, these results confirm that downregulation of NCoR1 and KLF16 results in activated mTOR signalling, resulting in decreased mucus (MUC2) synthesis and secretion.

**Fig 9.**
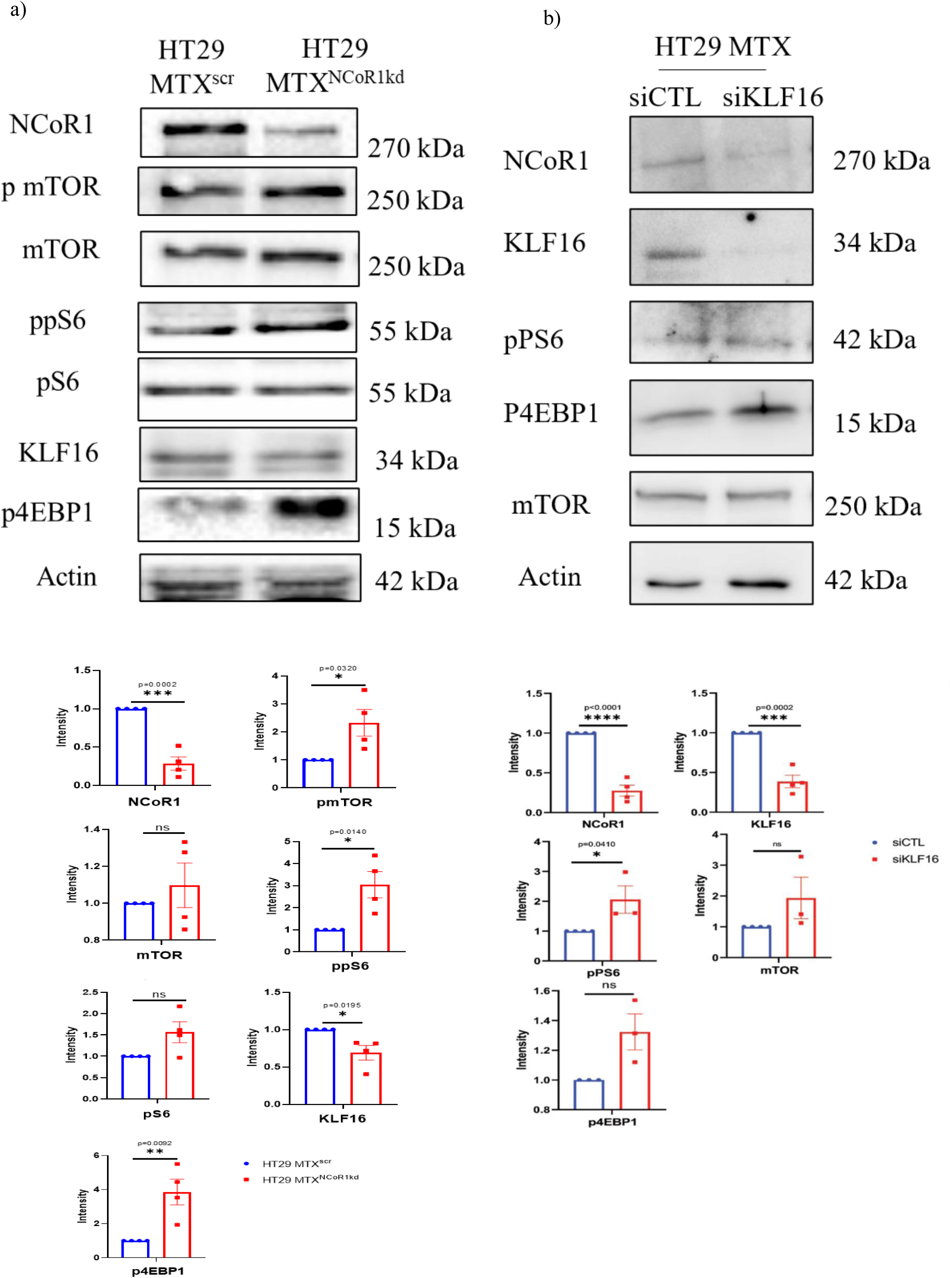
Activation of mTOR pathway upon NCoR1 and KLF16 perturbation. a) Immunoblot of NCoR1, KLF16 and mTOR pathway proteins pmTOR, mTOR, pS6, ppS6, p4EBP1, KLF16 and NCoR1 in the cell lysates of HT29 MTX ^NCoR1kd^ andHT29 MTX^scr^ cells. Graph below represents densitometric analysis showing fold intensity of respective protein expression calculated by normalising to loading control (Actin). b) Immunoblot of NCoR1, KLF16 and mTOR pathway proteins ppS6 and p4EBP1in the cell lysates of siCTL and siKLF16 HT29 MTX cells. Graph represents densitometric analysis showing fold intensity of respective protein expression of NCoR1, KLF16, ppS6, mTOR, and p4EBP1, calculated by normalizing to loading control (Actin). Each dot represents (a and b) an individual experiment. All data were expressed as means ± SEM, unpaired Student t-test was used to calculate statistical significance \**P* < 0.05; \*\**P* < 0.01; \*\*\**P* < 0.001; \*\*\*\**P* < 0.0001; ns, not significant.

**Fig 10:** *Representative Image:* (created with BioRender.com) Nuclear Co-repressor1 or NCoR1 is predominantly present in the crypts of the colon in healthy conditions, but during ulcerative colitis, its expression is reduced in the crypts, and its expression increases in the infiltrated immune cells. Due to the varied expression of NCoR1, functions of crypt residential cells alter during the onset of inflammation. In the goblet cell, a balanced regulation of NCoR1 and its target gene KLF16 is maintained, pursuing a regulated mTOR pathway and a regulated mucus secretion is attained during healthy conditions. But, upon inflammation, loss of NCoR1 and KLF16 in the goblet cell tends to activate the mTOR pathway. This upregulated mTOR pathway leads to reduced mucin production, affecting the goblet cell function during UC pathogenesis.

## Discussion

In recent developments modulating IBD Pathogenesis, molecular mechanisms driven by hyperactive immune regulation have been well reported. A constant epithelial-immune cell crosstalk is required for gut homeostasis via a proper mucosa. Impaired mucus barrier and epithelium exacerbate inflammation due to infiltrated microbiome-mediated dysbiosis, eventually leading to immune recruitment triggering inflammation^4^. Understanding the goblet cell mechanism that regulates mucus homeostasis, particularly during ulcerative colitis pathogenesis^30,31^, becomes important. In this study, we identify a distinct role for NCoR1 in maintaining mucosal barrier integrity by regulating crypts and goblet cells in the colon, which was previously known to regulate the immune cells^32-33^. The role of NCoR1 in goblet cell regulation has not yet been reported. Here, we highlight the importance of crypt residential NCoR1 in maintaining the goblet cells and preventing inflammation.

NCoR1 is classically characterized as a transcriptional repressor known to repress genes via recruitment of histone deacetylases^13^. The role of NCoR1 in combating intestinal inflammation is not well- understood^15^. NCoR1 modulates several pathophysiological conditions^34,13^, indicating its diverse functions. Our study reveals a dynamic expression of NCoR1 and its altered spatial distribution in UC. The expression patterns are dynamically altered upon disease severity. Crypt resident NCoR1 gets downregulated, and immune cells’ specific NCoR1 gets upregulated during inflammation. The shift in the expression pattern of NCoR1 tends to mark a clear indication of colitis.

Since UC majorly affects the mucosa, understanding the epithelial cell signaling becomes important. Loss of NCoR1 in the crypts becomes a major concern. Mucus barrier and goblet cell dysfunctions are widely reported during IBD^9,35^. Expression of NCoR1 was seen to be upregulated in goblet cells, indicating the potential role of NCoR1 in goblet cell function. Stable knockdown of NCoR1 in HT29 cells upon differentiation revealed an attenuated goblet cell phenotype. This suggests a very important role of NCoR1 in goblet cell maturation and its function in MUC2 formation. NCoR1 could be inhibited by targeting global HDACs using TSA, which is involved in the chelation of zinc cations from the active sites of HDACs, inhibiting recruitment of the NCoR1 complex, hence downregulating NCoR1 expression^26,27^. NCoR1 was upregulated via the induction of vitamin D3, as NCoR1 is recruited by the induction of VDR^36^. The exact modulation of NCoR1 was observed in the goblet cell culture.

To understand the mechanistic details *in vivo*, in the current work, we also generated a morpholino-based NCoR1 knockdown and Vitamin D-mediated NCoR1 overexpressed mouse model system. Knockdown of NCoR1 was observed at RNA levels. Loss of NCoR1 led to reduced goblet cell number and exacerbated inflammation, indicating its role in mediating goblet cell homeostasis. Vitamin D supplementation is an important medication to combat IBD, by an unspecified mechanism^37,38^. Here, we cite one of the perspectives that could be achieved by vitamin D induction, i.e., rescuing NCoR1 in the crypts, which in turn rescues the goblet cell number during inflammation. Overall, an important aspect of NCoR1-mediated crypt residential goblet cell homeostasis has been highlighted here.

NCoR1 complex-mediated gene repression involves HDACs ^39^. NCoR1-bound DNA were identified via ChIP sequencing, and NCoR1-bound genes were shortlisted based on the potential to modulate goblet cell function. Of several targets, KLF16 was observed to show higher binding to NCoR1 in differentiated goblet cells, marking its importance in cellular regulation. KLF16, a transcription regulator, has been identified here as a critical target of NCoR1 in regulating goblet cells. Here, we observe correlated expression of NCoR1 and KLF16, a phenomenon that contradicts the actual mode of working of NCoR1. However, the detailed insights on how NCoR1 regulates KLF16 need further understanding. We here consider KLF16 as an important target under the regulation of NCoR1 in the goblet cell. KLF16 is known to regulate endocrine activity in hepatic cells^40^, cortical neurons^41^ or uterine endometrial cells^42^ to maintain homeostasis. Regulation mediated by KLF16 is context-dependent, as a transcriptional activator or repressor^42,43^. Here, we observe that downregulated KLF16 decreased MUC2 and NCoR1, hence affecting the goblet cell function, exhibiting a co-regulatory mode of action along with NCoR1. KLF16 is observed to be downregulated during colitis. Knockdown of NCoR1 is seen as sufficient to downregulate KLF16 *in vivo,* indicating the importance of KLF16-mediated goblet cells and mucus regulation of the colon. The exact mechanism of how KLF16 is involved during inflammation and its potential to regulate NCoR1 needs further investigation. Also, cytokine or chemokine profiling has not been performed here, considering the mucus regulation as a primary function of the goblet cell. Though markers of inflammation *in vivo* were visibly observed and have also been previously reported^1,4^.

Several cellular signaling pathways are seen to be altered during IBD^2,44^. Knockdown of NCoR1 and/or KLF16 alone is seen to activate the mTOR pathway. Loss of NCoR1 and KLF16 upregulates phosphorylated forms of p-mTOR, p-pS6 kinase and p-4E-BP1, indicating the activation of the mTOR pathway. pS6 is primarily involved in cellular differentiation^45^. 4E-BP1 regulation is involved in cell growth and protein synthesis^46^. The activated mTOR pathway also contributes towards curtailment in mucin MUC2 production in goblet cells^47^. Activation of the mTOR pathway exacerbates IBD. Targeting the mTOR pathway as a therapeutic to combat inflammation has already been reported^28,29,48^. We hereby report the importance of the mTOR pathway with respect to goblet cell function. However, the phenomenon of mTOR activation during IBD is possibly multifactorial.

**Fig 10:**
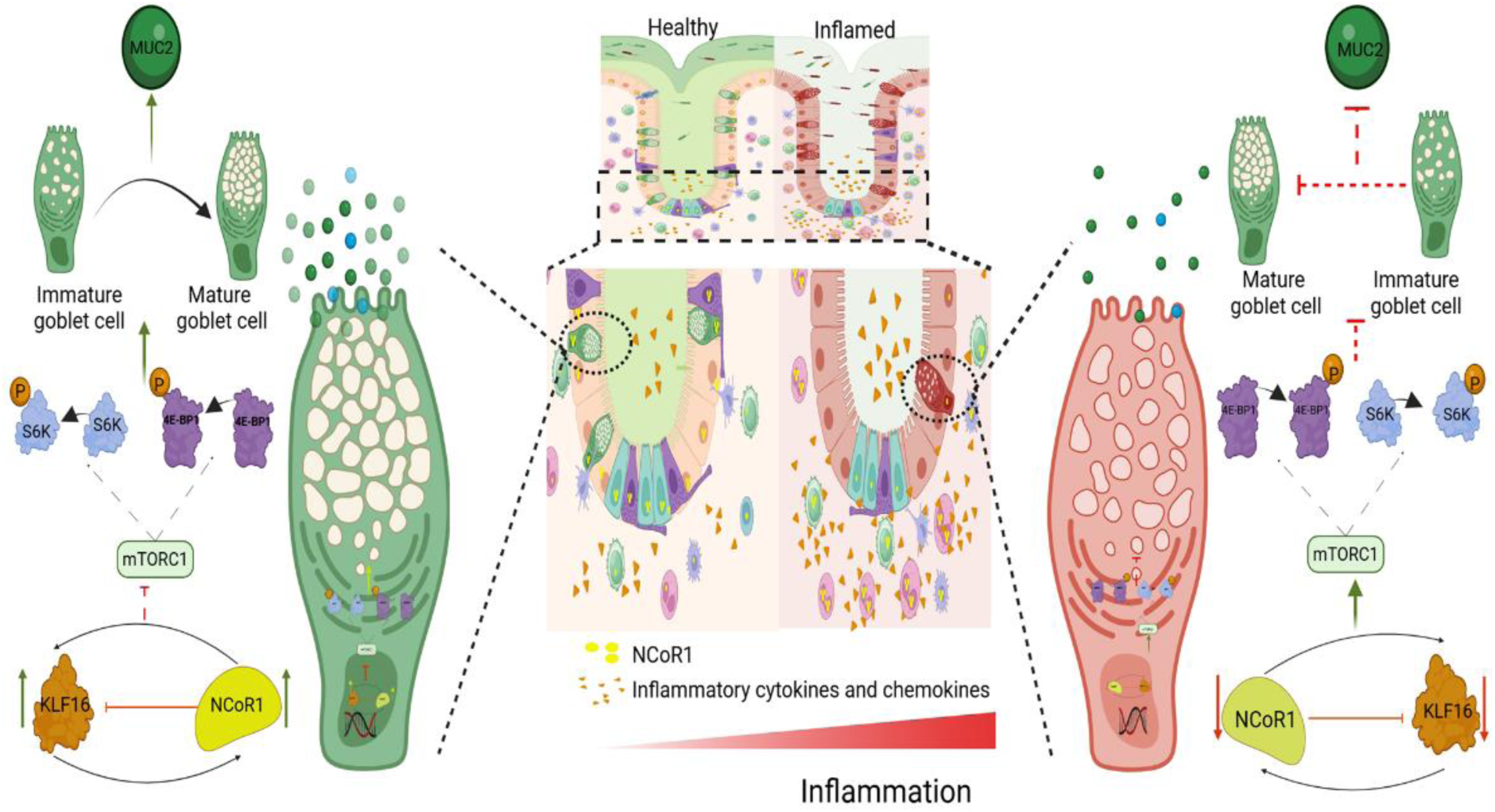
Graphical summary demonstrating the role of NCoR1-KLF16 axis regulating MUC2 in goblet cells via mTOR signaling pathway alteration.

## Materials and Method

### 1. Human colon biopsy samples

Human samples were collected from the Department of Gastroenterology at the All India Institute of Medical Sciences (AIIMS), New Delhi, in accordance with the standard biopsy collection protocol and with the required ethical clearance. These included Non-IBD (IBD suspects) (n=33) and UC(n=46) groups with ages between 18 and 60 years, with inclusion and exclusion criteria specified^49^. Two to four biopsies from each human subject were collected for qRT-PCR, Western blotting, and sectioning. Informed consent forms were acquired from all the patients. The influence of gender was not considered in the present study. UC severity was determined based on the ulcerative colitis endoscopic index of severity (UCEIS). Ethics approval for the use of human samples was obtained from both the institutes, the Regional Centre for Biotechnology (RCB) (IEC-141/05.03.2021, RP-34/2021) and AIIMS (IEC/NP-189/2013&RP-12/17.06.201307.06.2013).

### 2. Animal Strains and colitis models

The C57BL/6 mouse strain was used for the study. Mice were bred and housed in pathogen-free conditions and provided with sterilized food and water at 25°C with a 12-hour light and 12 hrs dark cycle in the Small Animal Facility. For inducing colitis, female C57BL/6 mice, 6-8 weeks old, weighing 18g-20g were used. Mice were fed with 2.5% DSS (w/v) for 3, 5 and 7 days in autoclaved water. Only autoclaved water was given to control mice. To knock down NCoR1in vivo, Vivo-morpholino was used; animals were given NCoR1-specific Vivo-morpholino and Control nontarget Morpholino. Two doses of intraperitoneal injections were given on the first day and third day during the course of DSS treatment at the concentration of 10 mg/kg (20nM/mice). Different conditions included in the study were: (i) Ctrl- mice were given non-target morpholino, (ii) Ctrl^NCoR1KD^ mice treated with NCoR1 Vivo morpholino, (iii) DSS mice along with non-target morpholino and (iv) DSS^NCoR1KD^ NCoR1 knockdown mice along with DSS treatment for 7 days. For upregulation of vitamin D-mediated NCoR1 in C57BL/6 mice, animals were fed with vitamin D3 at a dosage of 3000IU/kg/day with 1% CaCl2 mixed with feed for 3 weeks (Ctrl^VD^). Animals were fed with 2% DSS (w/v) for 1 week (3rd week) (DSS). A group of animals were treated with both vitamin D throughout 3 weeks, along with 2% DSS (only for 3rd week of the experiment) (DSS^VD^), called the rescue group. Body weights were monitored daily along with colitis symptoms like rectal bleeding and diarrhea. Colon and spleen were harvested. Colon and spleen were examined for changes in length and size. Organs were further used for qRT-PCR and Western blots as required. Distal colons were fixed with 4% formaldehyde and processed for sectioning.

### 3. Crypt isolation

Frozen colon samples were thawed and cut into 2-3 pieces of 1-2cm each and washed in 15ml PBS once, followed by consecutive DPBS washes until the solution becomes clear. Colon pieces were washed in 15ml chelation buffer (2% sorbitol, 1%sucrose, 5%FBS in DPBS) once. Colon were incubated in chelation buffer with 10mM EDTA for 20 mins under constant shaking at 4°C. The crypts were vortexed for 30 seconds, followed by 30 sec rest for five times. The supernatant was filtered using 100µm filter and collected, and the remaining crypts were scraped to detach them and filtered into the respective tubes. The tubes were centrifuged at 800g for 10 mins at 4 degrees to collect the crypts. The crypts were washed in PBS and collected in 2X Lamillie buffer and denatured at 95oC for 10 mins and quantified further using the CBX kit. Lysates were subjected to Western blot.

### 4. Histology and Immunostaining

Formalin-Fixed Paraffin-Embedded tissue were deparaffinized and were subjected to Hematoxylin-Eosin staining or Alcian blue–Nuclear Fast red staining as per requirement. Post staining, the slides were processed for dehydration steps via gradient ethanol washes followed by xylene wash. The slides were then mounted with coverslips after drying using DPX mountant. Sections were visualized in a Nikon bright-field microscope, and mucus thickness was measured using ImageJ software.

### 5. Immunohistochemistry

For immunostaining, human biopsy and mice colon samples were sectioned and after fixing with 4% paraformaldehyde antigen retrieval was performed. The sections were probed for NCoR1 (1:300) and incubated at 4 degrees overnight. The sections were probed with HRP tagged antibody for 2hrs followed by DAB staining. The sections were subjected to Alcian blue-Nuclear fast red staining. The slides were mounted using DPX mountant post drying. Sections were visualized in a Nikon bright-field microscope, and mucus thickness was measured using ImageJ software.

### 6. Cell culture and transfection

HT29 (Lot. 09K003) and HT29-MTX-E12 (Lot. 18K206) cell lines were obtained from ECACC. HT29 cells were cultured in DMEM medium. For differentiating into goblet cells, HT29 cells were grown in glucose-free DMEM, supplemented with galactose (250mM) for 3 days. HT29-MTX-E12 cells were grown in DMEM glutamax medium. Plasmid transfections and siRNA-mediated knockdowns were done along with cell seeding using Transfectin and Dharmafect, respectively. Required Plasmids and siRNA were incubated with the transfection reagent for 15 minutes in OptiMem media and further added to wells with cells. Stable NCoR1 knockdowns were created using NCoR1-specific shRNA via lentiviral transfection method using CalPhosTM Mammalian Transfection Kit, following the manufacturer’s protocol. 25µM vitamin D3 or 5µM tricostatin A treatments were given to cells for 24 hours. Cells were processed for ChIP seq as per protocol^50^. In house qRT-PCR were performed using gene specific primers and thermocycler conditions.

### 7. Immunocytochemistry

Cells seeded on 18mm glass coverslips were fixed with 4% methanol free paraformaldehyde and blocked in 0.1% BSA+0.01% TritonX-100 for 1 hour. Cells were probed with anti-Muc2 (1:400) and anti-NCoR1(1:300) at 4℃ overnight. Cells were further incubated with fluorophore-tagged secondary antibodies for 2 hours. Nucleic acid was stained with DAPI (1μg/ml). Coverslips were mounted with Prolong diamond Antifade Reagent and visualized in a confocal microscope (Leica SP8) for immunocytochemistry

### 8. Quantitative RT-PCR

Total RNA from the human biopsy samples, mice colon tissue or cultured cells was isolated using the RNeasy mini kit according to the manufacturer’s protocol. cDNA was prepared using 1μg of total RNA for each sample using a cDNA synthesis kit. Real-time PCR was performed utilising SYBR green master mix for 20μl reaction volume in 96-well plates in the CFX96 Real-Time system (BioRad). 18S, GAPDH or HPRT genes were used as housekeeping to normalize the reactions according to the sample origin.

### 9. Western blotting

Human biopsies and mouse organ tissues were homogenized using a tissue homogeniser (Precellys) in RIPA buffer to prepare protein lysates. Cells were lysed in RIPA buffer after PBS wash. Protein amount was measured by BCA assay. An equal amount of protein samples was separated on SDS-PAGE gel (4-12%) electrophoresis and transferred to nitrocellulose membrane. Blots were blocked with 5% skim milk for 1 hour at room temperature and probed with primary antibodies (1:1000) at 4℃ overnight against the desired proteins. Specific secondary antibodies (1:10,000) conjugated with HRP were probed for 1 hour at room temperature. Blots were detected and visualized for protein bands with Immobilon Forte western HRP substrate and imaged in Image Quant LAS4000. Band intensities were measured by ImageJ software.

### 10. Statistical Analysis

Results were analyzed and plotted using GraphPad Prism 8.0.1 software. All results were expressed as mean standard error from individual experiments done in triplicate. Data were analyzed with the standard unpaired Student’s t test and the Welch’s t test where applicable. Differences were considered significant at *P* values of \**p* < 0.05, \*\**p* < 0.01, \*\*\**p* < 0.001, and \*\*\*\**p* < 0.0001, as indicated in the figures. Error bar indicates means ±SEM.

---

**Fig S1:**
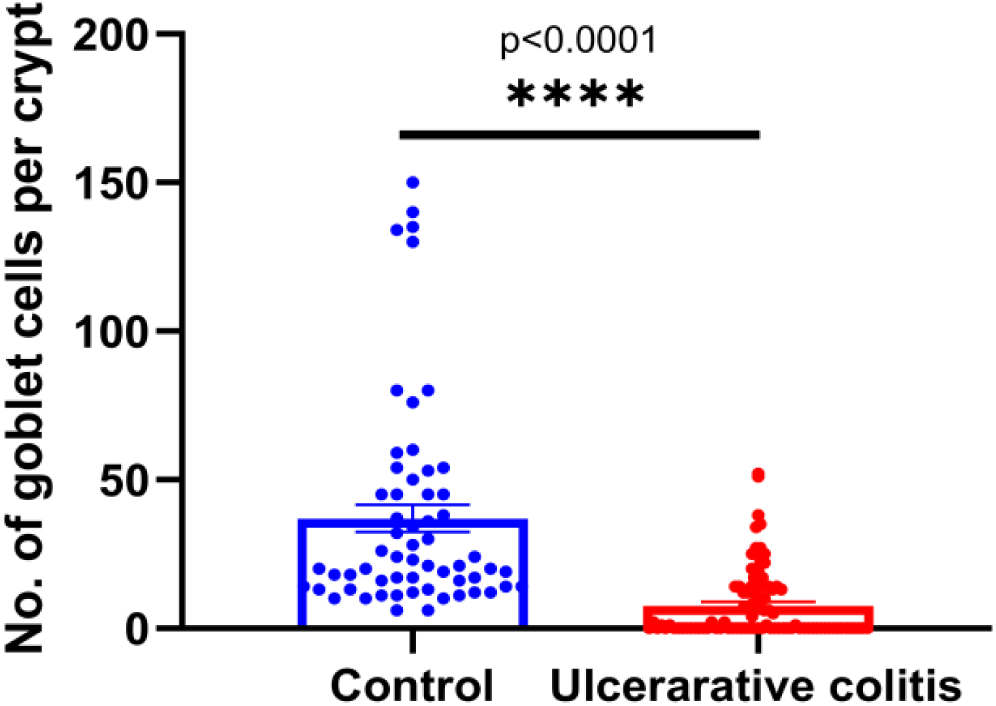
goblet cell number per crypt in control and UC patient. a) Graph showing goblet cell count in the crypts of human colon from control (n=60 crypts) and UC patients (n=93 crypts). All data were expressed as means ± SEM, unpaired Student t-test was used to calculate statistical significance \**P* < 0.05; \*\**P* < 0.01; \*\*\**P* < 0.001; \*\*\*\**P* < 0.0001; ns, not significant.

**Fig S2:**
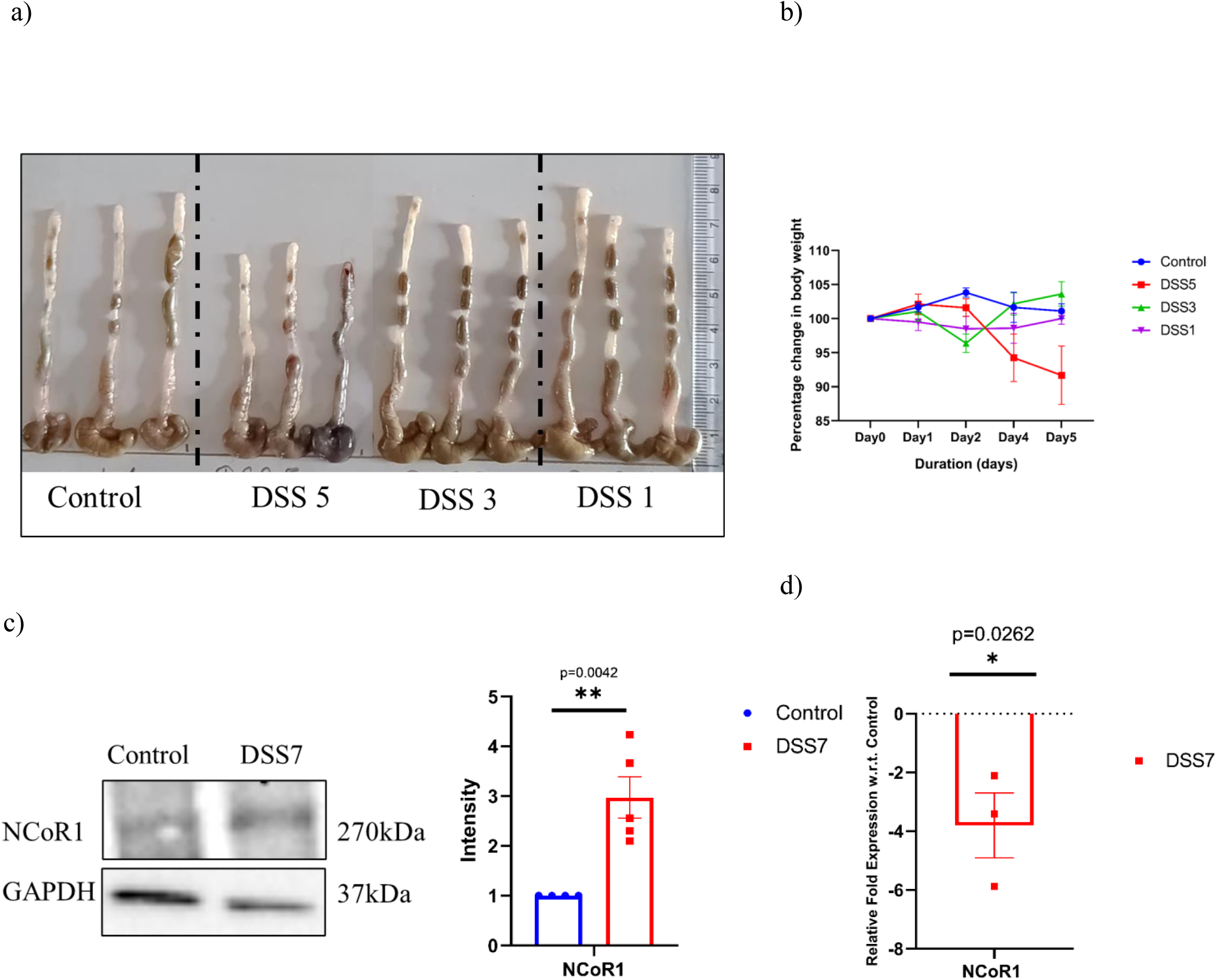
Expression of NCoR1 in acute murine colitis. a) Gross morphology of colon and caeca of control, DSS5, DSS3 and DSS1mice. b) Graph showing percentage body weight change of control, DSS5, DSS3 and DSS1mice. c) Representative Immunoblot of NCoR1 protein in whole tissue lysate of control and DSS7 mice (n=4 per group). The graph represents densitometric analysis showing fold intensity of NCoR1 expression calculated by normalizing to loading control (GAPDH). d) qRT -PCR analysis of relative fold expression of NCoR1 in colon tissues of DSS mice. GAPDH was used for normalization. Each dot represents (c and d) individual mice. All data were expressed as means ± SEM, unpaired Student t-test was used to calculate statistical significance \**P* < 0.05; \*\**P* < 0.01; \*\*\**P* < 0.001; \*\*\*\**P* < 0.0001; ns, not significant..

**Fig S3:**
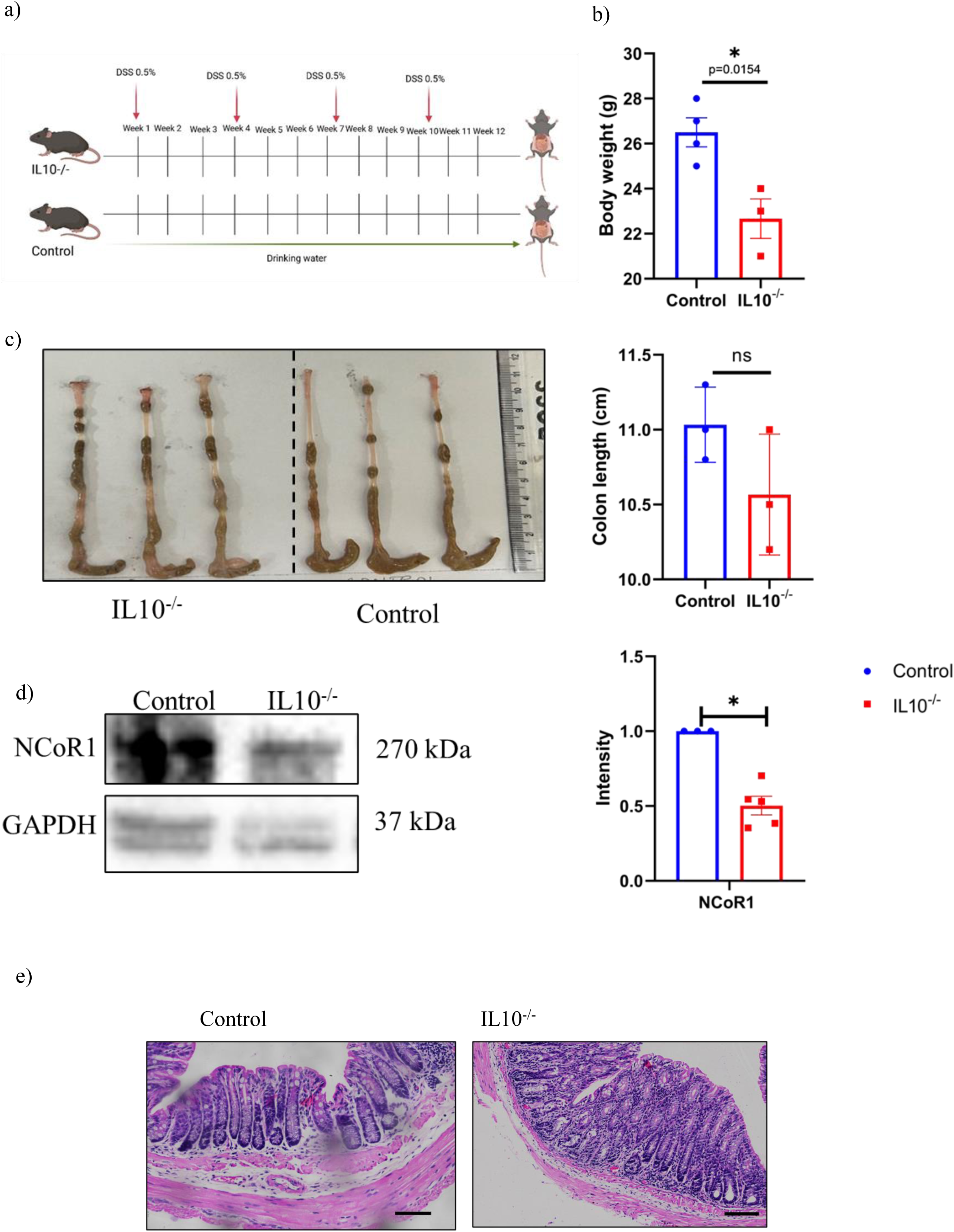
Expression of NCoR1 in IL10^-/-^ mice colitis. a) Schematic representation of the experimental plan for chronic colitis in IL10^-/-^mice using alternate DSS treatment (Point of administration indicated by red arrows) of 0.5% DSS in drinking water treatment for 12 weeks. (Created with BioRender.com) b) Graph showing the difference in body weight of control and IL10^-/^mice. c) Gross morphology of colon and caeca of control and IL10^-/-^ mice. The graph on the right shows colon length quantification. d) Representative immunoblot of NCoR1 protein in whole tissue lysate of Control and IL10^-/-^ mice. Graph represents densitometric analysis showing the intensity of NCoR1 expression calculated by normalizing to loading control (GAPDH). Each dot represents (b, c and d) individual mice. All data were expressed as means ± SEM, unpaired Student t-test was used to calculate statistical significance \**P* < 0.05; \*\**P* < 0.01; \*\*\**P* < 0.001; \*\*\*\**P* < 0.0001; ns, not significant. e) Haematoxylin Eosin staining in the distal colon tissue sections of control and IL10^-/-^mice (n=3 per group) (scale bar= 50um).

**Fig S4:**
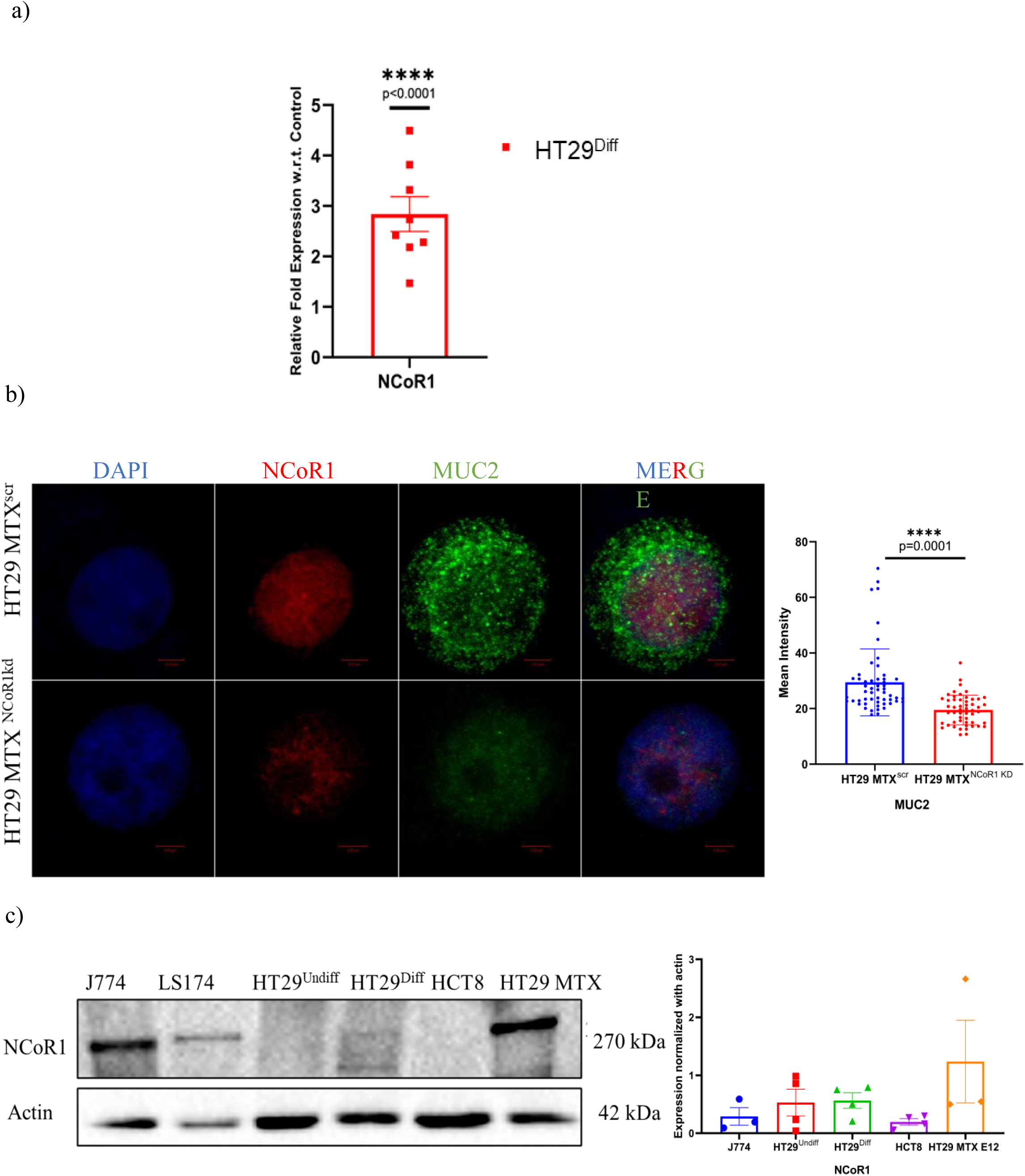
NCoR1 mediated alterations in HT29 and HT29 MTX goblet cells. a) qRT-PCR analysis of relative fold expression of NCoR1 genes in HT29^Diff^ cells, with respect to control HT29^undiff^ cells. HPRT was used for normalization. b) Confocal imaging of MUC2 (green) stained using anti-MUC2 and NCoR1 (red) stained using anti-NCoR1 antibodies in HT29 MTX ^NCoR1kd^ andHT29 MTX^scr^. (scale bar=3um). Graphs on the right show MUC2 fluorescence intensity measured. c) Immunoblot of NCoR1expression in J774, HT29^Undiff^,HT29^Diff^, HCT8 and HT29-MTX cell lines. Graph represents NCoR1 expression calculated by normalizing to loading control (Actin) in respective cell line. Each dot represents (a,b and c) individual experiment. All data were expressed as means ± SEM, unpaired Student t-test was used to calculate statistical significance \**P* < 0.05; \*\**P* < 0.01; \*\*\**P* < 0.001; \*\*\*\**P* < 0.0001; ns, not significant.

**Fig S5:**
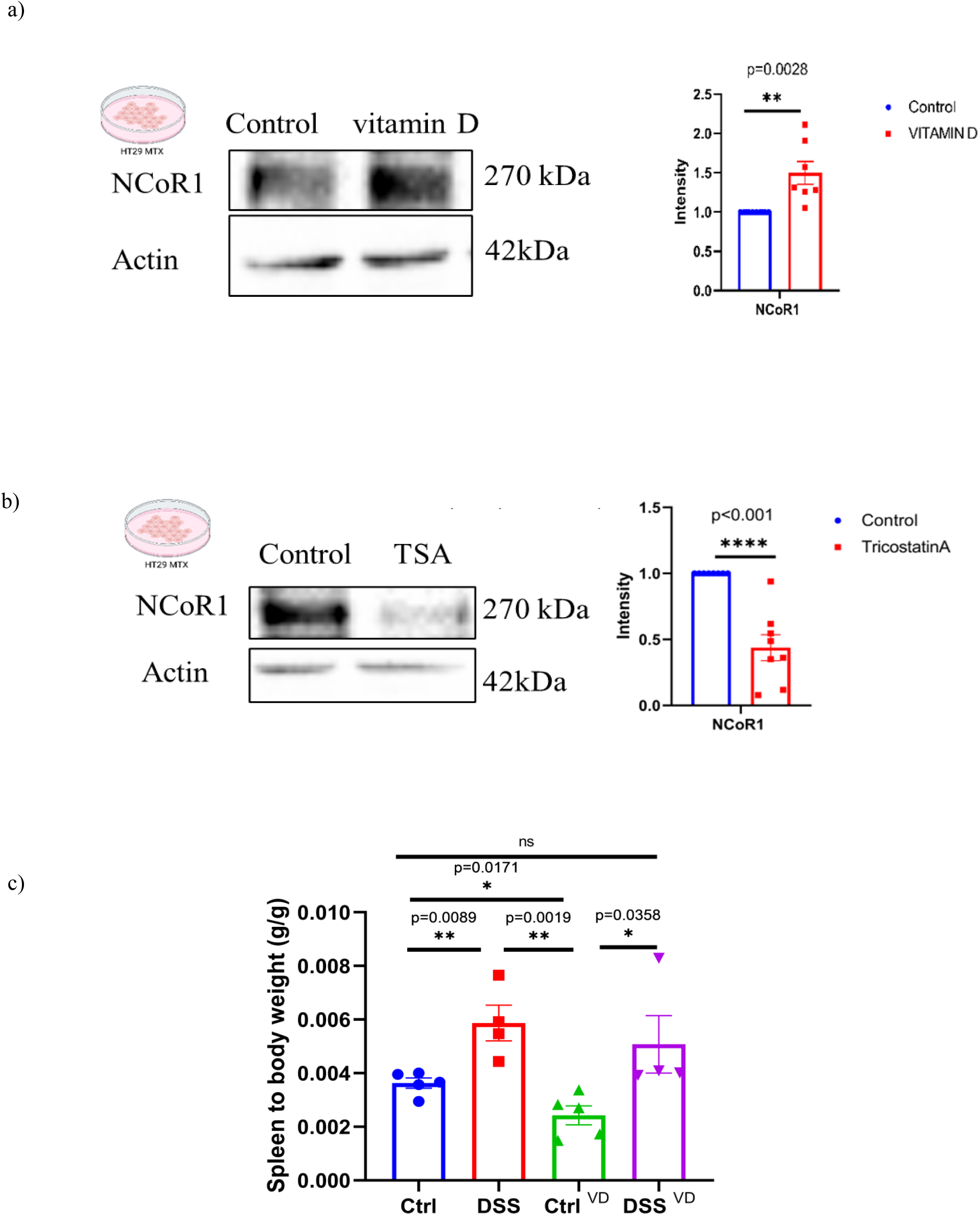
Perturbation of NCoR1 via vitamin D and Tricostatin A. a) Immunoblotting of NCoR1 in HT29 MTX cells upon vitamin D treatment at a concentration of 25uM for 24 hours. The graph on right represents densitometric analysis showing fold intensity of NCoR1 expression calculated by normalising to loading control (Actin) b) Immunoblotting of NCoR1 in HT29 MTX cells upon Tricostatin A treatment at a concentration of 5uM for 24 hours. The graph on right represents densitometric analysis showing fold intensity of NCoR1 expression calculated by normalising to loading control (Actin) c) Graph showing spleen to body weight ratio in Ctrl, DSS, Ctrl VD or DSS VD mice. Each dot represents (a and b) individual experiment and (c) individual mice. All data were expressed as means ± SEM, unpaired Student t-test was used to calculate statistical significance \**P* < 0.05; \*\**P* < 0.01; \*\*\**P* < 0.001; \*\*\*\**P* < 0.0001; ns, not significant.

**Fig S6:**
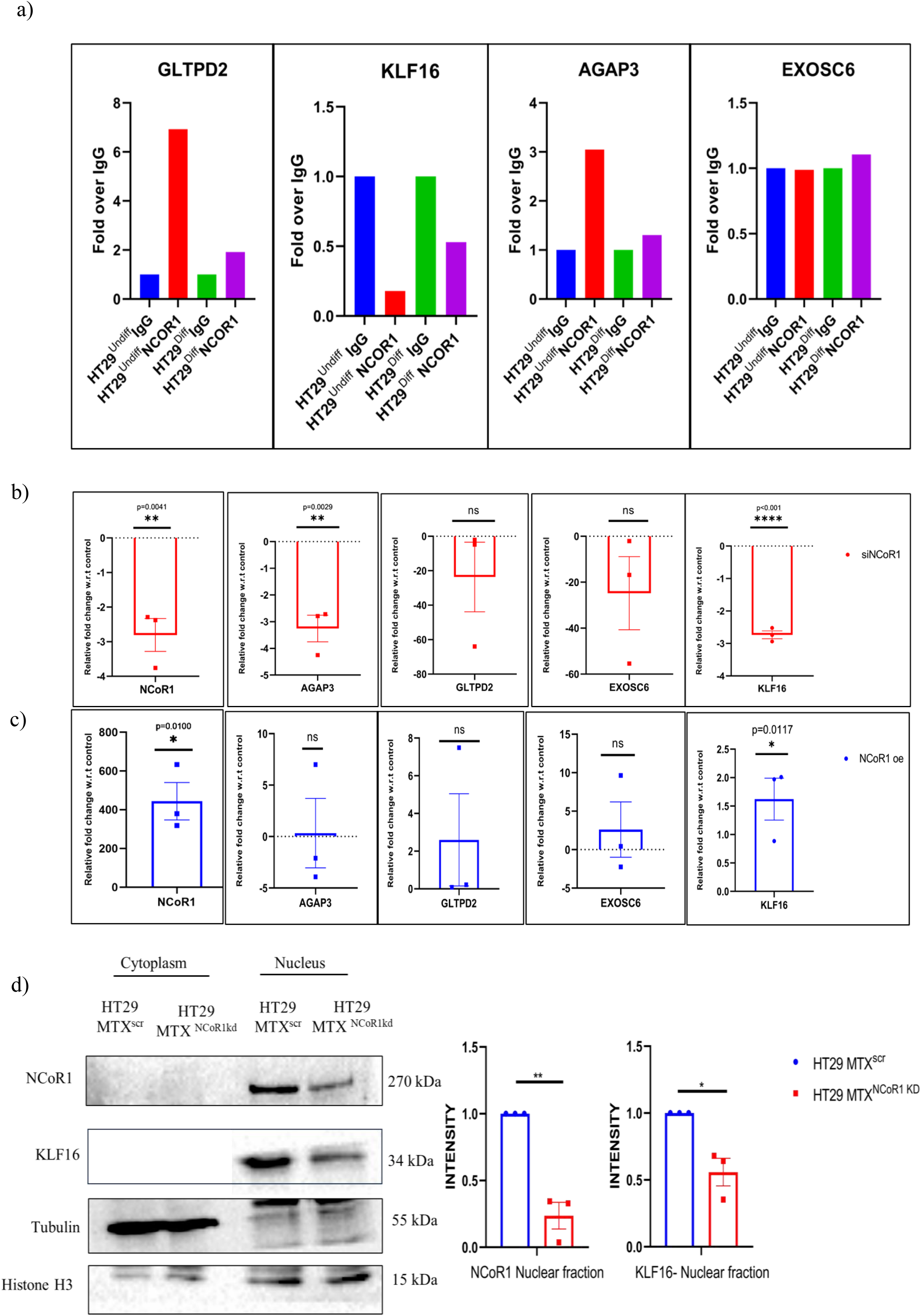
NCoR1 mediated regulation of target genes in goblet cell upon NCoR1 perturbation. a) Graph showing binding of AGAP3, KLF16, GLTPD2 and EXOSC6 in chromatin immunoprecipitation (ChIP) using anti- NCoR1 and anti-IgG antibodies in HT29^Undiff^, HT29^Diff^ cells. quantification done using fold over IgG method. b) qRT-PCR analysis of relative fold expression of NCoR1, AGAP3, GLTPD3, EXOSC6 and KLF16 genes upon transient knockdown of NCoR1 with respect to control HT29 MTX cells. GAPDH used for normalization. c) qRT-PCR analysis of relative fold expression of NCoR1, AGAP3, GLTPD3, EXOSC6 and KLF16 genes upon transient overexpression of NCoR1 with respect to control HT29 MTX cells. GAPDH used for normalization. d) Immunoblot of NCoR1 and KLF16 in Nuclear and cytoplasmic fractions of HT29 MTX^scr^ and HT29 MTX ^NCoR1kd^ cells. Graph on the right represents densitometric analysis showing fold intensity of NCoR1 and KLF16 expression in the nuclear fraction, calculated by normalizing to loading control (H3 for nuclear fraction). All data were expressed as means ± SEM, unpaired Student t-test was used to calculate statistical significance \**P* < 0.05; \*\**P* < 0.01; \*\*\**P* < 0.001; \*\*\*\**P* < 0.0001; ns, not significant.

**Fig S7:**
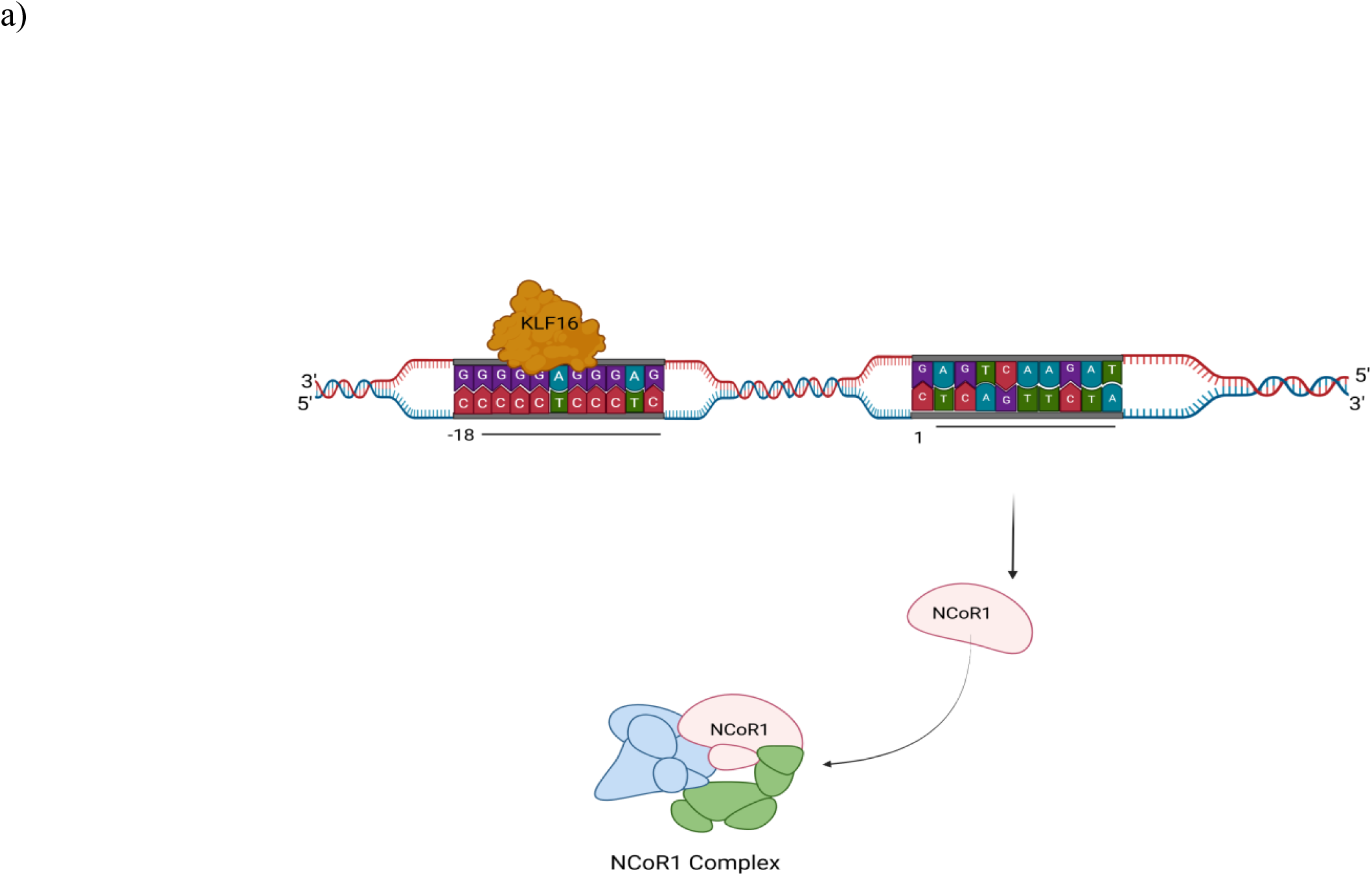
Promoter analysis of NCoR1. Image showing pSCAN predicted site for KLF16 binding on NCoR1promoter. Image created with BioRender.com

## Acknowledgment

We thank the sophisticated instrumentation facility at the Central Instrumentation Facility (CIF) of the Regional Centre for Biotechnology (RCB), the Flow Cytometry, Confocal Microscopy, Genomics, and Mass Spectrometry facility of the Advanced Technology Platform Centre (ATPC) in the RCB campus, and the Experimental Animal Facility (EAF) at NCR-Biotech Science Cluster for animal experiments. We are thankful to Professor Johan AUWERX (EPFL, Switzerland) for FLAG-NCoR1 plasmid. We thank Chromosome Labs Pvt Ltd (Mumbai, India) for ChIP sequencing Data generation and analysis. We thank Professor Manjula Kalia (RCB), Professor Avinash Bajaj (RCB), Dr. Pallavi Kshetrapal (associate professor, TSHTI) for their extended support and Inputs.

## Author Contributions

Conceptualization: Chittur V. Srikanth.

Methodology: Yesheswini Rajendran, Bhagyashree Srivastava and Chittur V. Srikanth.

Investigation: Yesheswini Rajendran, Bhagyashree Srivastava, Muskaan Kalra, Neha Guliya, Rohan Babar.

Resource: Preksha Gaur, Aamir Suhail, Mukesh Singh, Lalita Mehra. Supervision: Prasenjit Das, Vineet Ahuja, and Chittur V. Srikanth.

Project Administration: Chittur V. Srikanth.

Writing-original draft: Yesheswini Rajendran and Bhagyashree Srivastava. Writing -review and editing: Yesheswini Rajendran and Chittur V. Srikanth. Animal ethics: RCB/IAEC/2024/101, RCB/IAEC/2023/177, RCB/IAEC/2024/201

Human Ethics :(RCB) (IEC-141/05.03.2021, RP-34/2021) and AIIMS (IEC/NP-189/2013&RP 12/17.06.201307.06.2013). (RCB)(RCB-IEC-H-11) and (AIIMS)(IEC-217/05.04.2019, RP-19/2019)

